# *Arabidopsis thaliana* Phytochrome A Sensory Properties in Canopy Shade

**DOI:** 10.1101/2025.05.14.654025

**Authors:** Philip Butlin, Marissa Valdivia-Cabrera, Ramon Grima, Karen J. Halliday

**Affiliations:** School of Biological Sciences, University of Edinburgh, Edinburgh 3H9 3BF, United Kingdom

**Author notes:** Authors have made equal contributions (Philip Butlin, Marissa Valdivia-Cabrera).

## Abstract

Canopy shade environments can negatively impact plant growth and survival. Phytochrome A (phyA) is essential for seedling adaptation to deep shade, yet its role under moderate natural canopy conditions remains unclear. By expressing a novel phyA-nanoLUC reporter in *Arabidopsis thaliana*, we quantify diurnal fluctuations in phyA and its induction by low red to far-red (R:FR) ratio, simulating canopy shade. We uncover a new key regulatory function for phyB, which increases phyA stability post-dawn in unshaded conditions and modulates expression of *PHYA* in low R:FR. Further, we demonstrate that beyond its role as a deep shade sensor, phyA is an adept sensor of canopy shade, capable of detecting a range of R:FR ratios across light intensities. Mathematical modelling demonstrates that this property arises from the dynamic features of the phyA HIR mode of action. Interestingly, phyA synthesis is strongly induced by subtle reductions in R:FR ratio, and is robust to light perturbation, suggesting the phyA-sensory module is configured to detect modest shade. Spectral data from natural shade habitats provides ecological context for our laboratory findings. Physiological analysis indicates that under canopy shade phyA promotes seedling de-etiolation, organises resource management and accelerates reproductive development. We propose that this suite of responses, termed the “canopy adaptation strategy”, enhances survival chances under conditions where shade-avoidance strategies are maladaptive.

## Introduction

In natural ecosystems, vegetational shade presents a critical challenge for plants by reducing the light available for photosynthesis and detrimentally impacting growth. Remarkably, many plants are not only capable of detecting when they are in shade, but also differentiating between types of shade. Crucially, this ability to distinguish between shade environments produced, for example, by encroaching neighbours directly competing for available light and those cast by overstory canopies, enables plants to deploy adaptive growth strategies tailored to the specific scenario they face (1, 2).

At the heart of this shade detection system are the phytochrome photoreceptors. Phytochrome B (phyB) plays a pivotal role in eliciting the shade avoidance syndrome (SAS), which includes hypocotyl elongation in seedlings, as well as leaf hyponasty, petiole elongation, reduced leaf area and accelerated flowering in adult plants (1, 3). By directing growth upwards and outwards, the SAS provides a competitive advantage to plants by improving leaf positioning and enhancing light interception amongst similarly sized neighbours (3–5). Conversely, adopting the SAS is considered to be a detrimental strategy in the most severe, ‘deep shade’ environments (e.g. in thick undergrowth), where excessive elongation growth reduces survivability (6). Plants, therefore, utilise a second phytochrome in deep shade, phyA, to antagonise the SAS in developing seedlings (6–8). While the importance of phyA action in deep shade is well documented (6–8), the efficacy of phyA beneath milder vegetational canopies, produced either by tree cover or ground-level overtopping neighbours and frequently encountered in natural environments, remains uncertain. Interestingly, as with deep shade scenarios, the adaptive value of shade-induced elongation in some species, including rosette annuals like *Arabidopsis thaliana*, is believed to diminish beneath canopy cover (5, 9).

Light in canopy shade is both reduced in photosynthetically active radiation (PAR; 400-700 nm) and spectrally enriched with far-red (FR; 700-780 nm), relative to red (R; 600-700 nm), wavelengths (4, 10, 11). The degree of reduction in the R:FR ratio can be used to quantify the severity of foliar shade (11). Changes in the external prevalence of R and FR wavelengths are monitored by phytochromes, a dichromic class of photoreceptors that exist in photoreversible inactive (Pr) and active (Pfr) forms (4, 12). R light is absorbed by Pr (λ_max_ ∼660 nm), which induces photoconversion to the active Pfr form. Conversely, FR absorption by Pfr (λ_max_ ∼730 nm) reverses this process, shifting phytochrome molecules to the inactive, Pr state (12). While all phytochromes share these photosensory traits, their ways of operating differ markedly. Most notably, phyA and phyB, the two most prominent of the five phytochromes (phyA-E) in Arabidopsis, have divergent response behaviours (13–15).

Modelling approaches have been instrumental in creating a theoretical framework for understanding the operational properties of phyA and phyB (16–22). PhyA functions in either a Very Low Fluence Response (VLFR) mode, induced by weak irradiances of any wavelength, or a High Irradiance Response (HIR) mode, activated by continuous irradiation with FR (14). The phyA signalling module includes a photocycle-coupled, nuclear shuttling mechanism involving homologous carrier proteins FAR-RED ELONGATED HYPOCOTYL 1 (FHY1) and FHY1-LIKE (FHL) (17, 23, 24). This process, driven by photoconversion between the Pr and Pfr forms, involves several steps. Initially, phyA-Pfr in the cytoplasm binds to FHY1/FHL, promoting its transport into the nucleus. As the signalling capacity of phyA-Pfr is impaired in this complex, two further photoconversions are required within the nucleus – first to Pr, releasing FHY1/FHL, and then back to Pfr – for uncoupled, biologically active Pfr to accumulate. This cyclical sequence, which is fundamental to the phyA response mechanism, helps explain the HIR fluence rate dependency and the distinctive FR-shift in its action spectra (17, 25).

In contrast, phyB operates in a Low Fluence Response (LFR) that is R/FR reversible. PhyB undergoes photoactivation in response to R light, resulting in translocation of phyB-Pfr from the cytoplasm to the nucleus to execute its signalling functions (16, 25). Absorption of FR irradiances, on the other hand, initiates the photoreversion of phyB-Pfr to -Pr, resulting in phyB nuclear expulsion (25). Model analysis has shown that the proportion of active (nuclear) phyB (Pfr/Ptot [Pr+Pfr]) under high irradiance light is strongly linked to the external R:FR (4, 22, 26, 27). This quality makes phyB effective in sensing reflected FR light from neighbouring plants where light levels are typically high (11, 22). However, in lower irradiance scenarios, such as those encountered in a closed-canopy environment or, more fleetingly, as a result of cloudy weather, thermal reversion of Pfr back to Pr becomes increasingly influential on Pfr/Ptot (22). Consequently, phyB activity being more strongly impacted by temperature in such scenarios and less so by R:FR (22). Interestingly, in Arabidopsis accessions such as *Landsberg erecta, Wassilewskija* or *Columbia* (studied here), no thermal reversion of phyA has been observed (28, 29). It is therefore worthwhile exploring the role of phyA in detecting canopy shade conditions where phyB may not serve as a reliable sensor of competition.

In this study, we set out to determine the operational range of phyA in canopy shade and how this relates to the unique properties of phyA HIR response mode. To do this, we developed phyA-nLUC lines that provide quantitative readout of phyA abundance. Our data show phyA levels are highly dynamic, responding to even very subtle reductions in R:FR ratio indicative of mild canopy shade. Mathematical modelling and experimental analysis established that the specialised photosensory properties of phyA allow it to accurately detect changes in R:FR ratio. As a result, phyA serves as a sensitive detector of canopy shade, triggering an adaptive growth strategy for canopy adaptation.

## Results

### Quantitative tracking of phyA abundance using phyA-nLUC reporters

PhyA protein abundance is highly dynamic, exhibiting strong diel rhythmicity (30, 31). To quantitatively track this dynamic behaviour we generated phyAp::phyA-nanoLUC (phyA-nLUC) constructs and stably expressed them in the *phyA-211* mutant background, successfully completing the phenotype (*SI Appendix,* Fig. S1). Our data show phyA-nLUC exhibits the expected daily rhythmicity in 10L:14D photoperiods, with phyA-nLUC accumulation at night and depletion during the day (Fig. 1 *A*; *SI Appendix,* Fig. S2 *A*). Consistent diurnal trends were observed across several reporter lines (*SI Appendix,* Fig. S3). This rhythmic behaviour is consistent with earlier findings that boh *PHYA* transcripts and phyA protein accumulate during the night, while phyA protein declines during the day due to the relative instability of phyA-Pfr (25, 28, 32). Rhythmicity in phyA levels was previously demonstrated in seedlings transferred from light-dark cycles to continuous light (CL) (30). We observed a low amplitude phyA-nLUC rhythm for 1-2 days following transfer to CL, but this rhythm was not sustained over time (*SI Appendix,* Fig. S2 *B* and S4). The phyA-nLUC signal depleted to 85% of ZT0 levels after 24 h in CL, and fell to a baseline of ∼36% by 110 h (*SI Appendix,* Fig. S2 *B*). Our data demonstrates that phyA-nLUC reliably reports phyA levels and offers a quantitative insight into its dynamics.

**Fig. 1.**
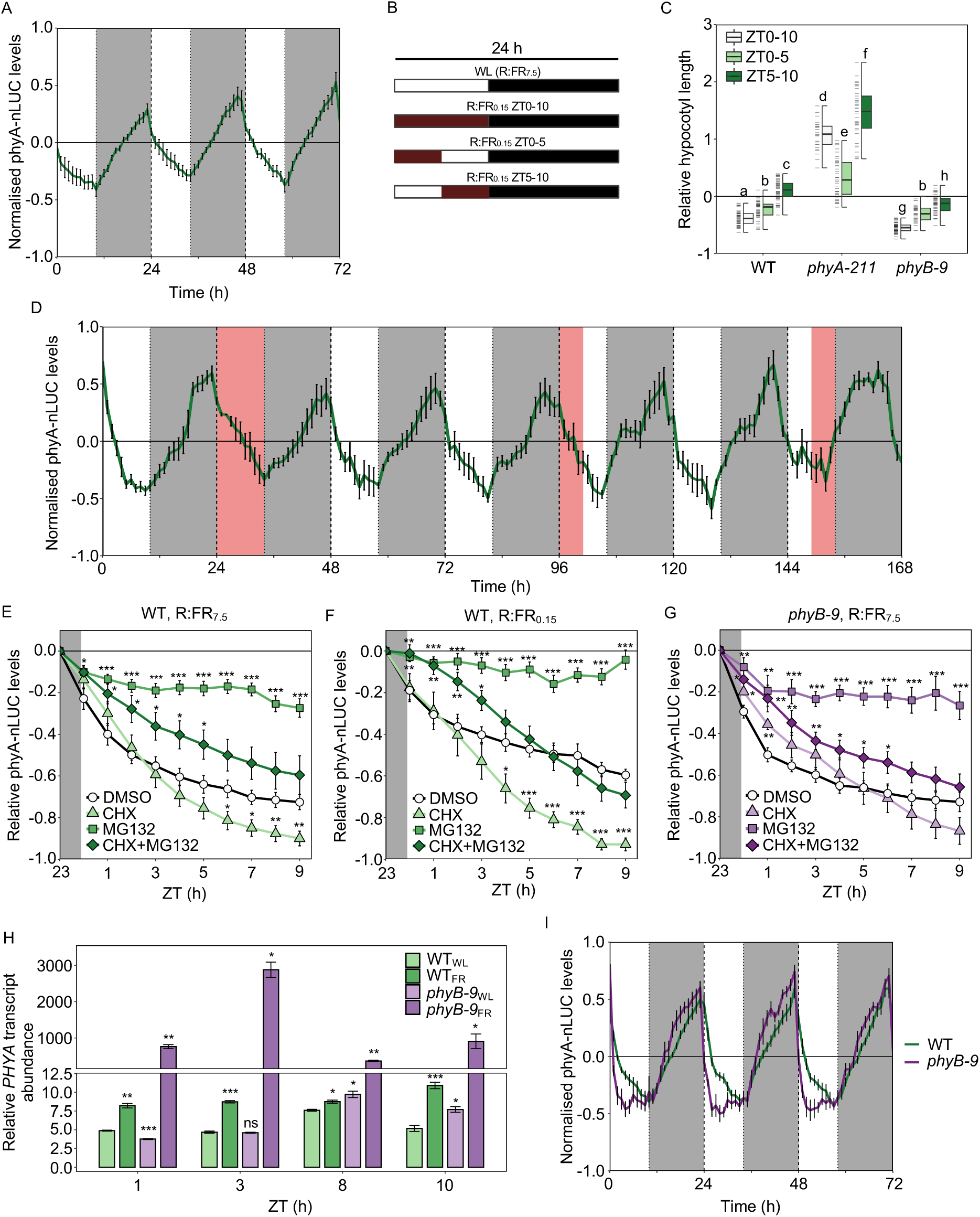
Low R:FR ratio light increases phyA abundance and activity at any time of day. (*A*) Normalised bioluminescence of phyA-nLUC in diurnal cycles (10L:14D) of white light (R:FR_7.5_). (*B*) Schematic of R:FR_0.15_ treatments (predicted Pfr/Ptot ∼0.24) provided in *C* and *D*. (*C*) Relative hypocotyl lengths (calculated with respect to the length of each genotype in R:FR_7.5_ [predicted Pfr/Ptot ∼0.80]) of 6-day-old WT, *phyA-211* and *phyB-9* seedlings treated daily with R:FR_0.15_ from ZT0-10, ZT0-5 or ZT5-10. (*D*) Normalised phyA-nLUC readings in entrained seedlings treated with R:FR_0.15_ between 24-34 h, 96-101 h or 149-154 h. Hourly abundance changes in phyA-nLUC expressed in WT (*E* and *F*) and (*G*) *phyB-9* seedlings treated with 50 μM MG132 and/or 200 μM CHX to block either proteasomal degradation, translation or both; DMSO alone was used as a control. Relative phyA-nLUC levels following the dark-light transition (ZT0) into a high (R:FR_7.5_; *E* and *G*) or low (R:FR_0.15_; *F*) day period, calculated with respect to the point of pre-dawn inhibitor application (ZT23). (*H*) Abundance of *PHYA* transcripts (normalised to *PP2A*) in WT or *phyB-9* seedlings at 4 time points throughout R:FR_7.5_ or R:FR_0.15_ day periods using qRT-PCR. (*I*) Normalised bioluminescence of phyA-nLUC in 10L:14D cycles in WT (green line) or a *phyB-9* (purple line) background. (*A*, *D*-*G* and *I*) Traces show mean signal produced by *n* ≥ 8 plants, measured at 1 h intervals; error bars show ± SEM. (*A*, *D* and *I*) Background colours of each panel correspond to R:FR_7.5_ (white), R:FR_0.15_ (red) and night (grey) periods. (*C*) Box plots display: the median (centre line); upper and lower quartiles (box limits); 1.5 x interquartile range (whiskers); individual data points (dashes on the left of boxes, *n* ≥ 25). Letters denote statistically indistinguishable groups according to a Kruskal-Wallis test followed by a *post-hoc* Dunn’s Test (Bonferroni correction). (*H*) Bars indicate the mean; error bars show ± SEM (*n* = 3). Asterisks indicate significant differences at each time point from DMSO (*E*-*G*) and WT_WL_ (*H*) controls (*\*P* < 0.05; *\*\*P* < 0.01; *\*\*\*P* < 0.001), calculated using a Student’s t-test. All experiments were conducted in a background WL of 15 μmol m^-2^ s^-1^. Each experiment was repeated three times.

### Delineating the contextual framework for phyA action

An initial aim of this study was to elucidate both the experimental and environmental parameters required to elicit phyA action in ‘canopy shade’ (i.e. overhead foliage produced either by an overstory or directly overtopping neighbours). To do this, we used the length of seedling hypocotyls as a quantitative, physiological measure of phytochrome activity (33). Preceding studies suggested that phyA action is prominent in deep shade (e.g. PAR ≤ 15 μmol m^-2^ s^-1^ and R:FR ≤ 0.15) during early seedling development (8, 34). Consistent with these reports, daytime low R:FR (R:FR_0.15_) treatments in 10L:14D photoperiods commencing on day 1 (D1), but not D3, effectively suppresses hypocotyl growth via phyA without impacting germination rates (*SI Appendix,* Fig. S5). All subsequent seedling experiments administered light treatments from D1 post-germination.

We previously established that phyA serves as a critical dawn sensor, primarily due to night-time phyA accumulation and photoactivation upon dawn light exposure (32). Further, the phyB shade-avoidance response is known to be circadian gated with peak sensitivity at dusk (35). To assess whether the sensitivity of the phyA activated response also varies with the time of day, we analysed hypocotyl growth under 10L (PAR = 15 μmol m^-2^ s^-1^, R:FR_7.5_):14D photoperiods when R:FR_0.15_ light was supplied in the morning (ZT0-5), afternoon (ZT5-10) or continuously through the 10-hour daylight period (ZT0-10; as illustrated by the schema in Fig. 1 *B*). We found that application of R:FR_0.15_ to *phyA-211* seedlings between ZT0-10 or ZT5-10 significantly promoted hypocotyl elongation relative to the R:FR_7.5_ control (Fig. 1 *C*). Therefore, as anticipated, eliminating the inhibitory effect of phyA resulted in enhanced growth during times that coincide with the evening peak sensitivity for phyB-mediated shade avoidance (35–37). In contrast, all R:FR_0.15_ treatments, suppressed hypocotyl extension in a phyA-dependent manner in WT, with similar effects observed in *phyB-9* (Fig. 1 *C*). We also found R:FR_0.15_ supplied during ZT0-10, ZT0-5 or ZT5-10 elevated levels of both phyA-nLUC (Fig. 1 *D*) and *PHYAp::LUC* (*SI Appendix,* Fig. S6 *A*), which reports *PHYA* promoter activity. Quantitative phyA-nLUC values for control vs R:FR_0.15_ (ZT0-10) are shown for ZT3, ZT6 and ZT9 replicate time points (*SI Appendix,* Fig. S6 *B*). Therefore, unlike the phyB-SAS, which displays a rhythmic response sensitivity that peaks in the evening (35), exposure to prolonged low R:FR periods can increase phyA production and activity at any time of day. Although phyA levels being low and mainly arrhythmic in continuous light (CL) (*SI Appendix*, Fig. S2 *B* and S4), we also found that R:FR_0.15_ was effective in elevating phyA-nLUC levels and suppressing hypocotyl growth in these conditions (*SI Appendix*, Fig. S4, S7). R:FR_0.15_ does, however, elicit significant hypocotyl elongation in CL, which obscures that phyA HIR response observed in photoperiodic (10L:14D) conditions (*SI Appendix*, Fig. S7).

### PhyA synthesis and destruction dynamics in canopy shade

Dynamic changes in phyA abundance are determined by both destruction- and synthesis-rates (25, 28, 30). To better understand the factors controlling phyA levels throughout the day, we investigated the effects of a proteasome inhibitor (MG132) and a protein synthesis inhibitor (cycloheximide; CHX) on phyA-nLUC levels. First, we observed that in a high R:FR (R:FR_7.5_) photoperiod, abundance of phyA-nLUC progressively declined towards a minima at ZT9 of 27% compared to pre-dawn levels (ZT23; Fig. 1 *E*). Application of MG132 effectively stemmed this daytime reduction in phyA-nLUC (declined to 73% of initial value by ZT9), deduced from the comparison of MG132 vs DMSO control, as well as MG132 + CHX (40% of ZT23 value) vs CHX alone (10% of ZT23 value; Fig. 1 *E*). Conversely, CHX treatment resulted in a significant decline in phyA-nLUC levels relative to DMSO or MG132, indicating that phyA is synthesised throughout the day (Fig. 1 *E*). Daytime phyA-nLUC abundance was more stable in R:FR_0.15_, declining to only 40% of pre-dawn levels (Fig. 1 *F*). Reducing the R:FR ratio, is expected to increase the proportion of the more stable Pr isoform, potentially reducing proteolysis (17, 19, 28). However, the impact of MG132 was slightly more pronounced in R:FR_0.15_ (daytime phyA-nLUC signal maintained on average at ∼ 92% of ZT23 level) compared to R:FR_7.5_ (∼ 82% of ZT23 level), indicating that proteolysis increases marginally in R:FR_0.15_ (Fig. 1 *E* and *F*). On the other hand, CHX application was more effective in reducing phyA-nLUC levels in R:FR_0.15_ (33% lower than DMSO at ZT9) compared to R:FR_7.5_ (17% lower than DMSO at ZT9). Thus, R:FR_0.15_ appears to elevate phyA-nLUC levels in 10 h photoperiods primarily through increased synthesis (Fig. 1 *F* and *SI Appendix,* Fig. S6 *A*).

### PhyB regulation of phyA dynamics

As there is evidence for phyA-phyB cross-talk (38–40), we next sought to assess the impacts of the *phyB-9* mutation on phyA-nLUC levels. Contrasting with the phyA-nLUC signal responses to R:FR_0.15_ in WT seedlings (Fig. 1 *F*), CHX had only a slight impact on daytime phyA-nLUC levels in *phyB-9* in R:FR_7.5_ (Fig. 1 *G*). Concurring with this, transcription analysis revealed that R:FR_0.15_ application was largely more effective than loss of phyB in inducing *PHYA* expression, although their effectiveness was more similar at ZT8 (Fig. 1 *H*). R:FR_0.15_ also had a more pronounced impact on *PHYA* in the *phyB-9* background, indicating that phyB plays a role in suppressing the phyA low R:FR response. In support of this, exposure to R:FR_0.15_ from ZT0-5 induces higher levels of phyA-nLUC in a *phyB-9* background (*SI Appendix*, Fig. S8) as well as a stronger hypocotyl suppression responses when R:FR_0.15_ is applied from ZT0-10 or ZT5-10 (Fig. 1 *C*).

We also found that phyA-nLUC accumulates at a faster rate at night in the *phyB-9* background (Fig. 1 *I* and *SI Appendix*, Fig. S9) and that lack of phyB significantly accelerates the decline of the phyA pool immediately following the dark-light transition (Fig. 1 *E*, *G* and *I*). Given that MG132 application blocks this decline (Fig. 1 *G*), the *phyB-9* mutation appears to increase proteasomal destruction of phyA. In summary, phyB appears to be a potent regulator of phyA, curbing low R:FR-induced *PHYA* expression, while enhancing phyA stability and daytime abundance in unshaded, high R:FR, photoperiods.

### PhyA is a reliable sensor of low R:FR ratio canopy shade

Given the complexity of the phyA signalling module and its fluence rate-dependency (7, 17), we next wanted to determine whether phyA can function as a dependable R:FR sensor in scenarios, such as canopy shade, where the R:FR detection capabilities of phyB breakdown (22). In order to considered a reliable R:FR sensor, we reasoned that phyA must be able to consistently respond to R:FR changes under a variety of light irradiances. We therefore quantified the phyA response across a range of R:FR ratios and at different R light (RL) fluence rates (8, 25, 50 or 100 μmol m^-2^ s^-1^). At RL_8_, we were able to deliver R:FR ratios of 1.5, 0.9, 0.6, 0.3 and 0.15, although limitations in FR capacity of growth chambers meant that the range of R:FR ratio treatments at higher RL fluence rates was more restricted (Fig. 2 *A*). Predicted Pfr/Ptot ratios for R:FR treatments are approximately: 0.86 (RL), 0.71 (R:FR_1.5_), 0.63 (R:FR_0.9_), 0.55 (R:FR_0.6_), 0.42 (R:FR_0.3_), and 0.26 (R:FR_0.15_) (*SI Appendix*, Fig. S10 *A* and Table S1). For data interpretation purposes, it is important to note that the capability of phyA to operate in a VLFR varies across Arabidopsis accessions and are not detectable in *Columbia*, the accession used in this study (37, 40).

**Fig. 2.**
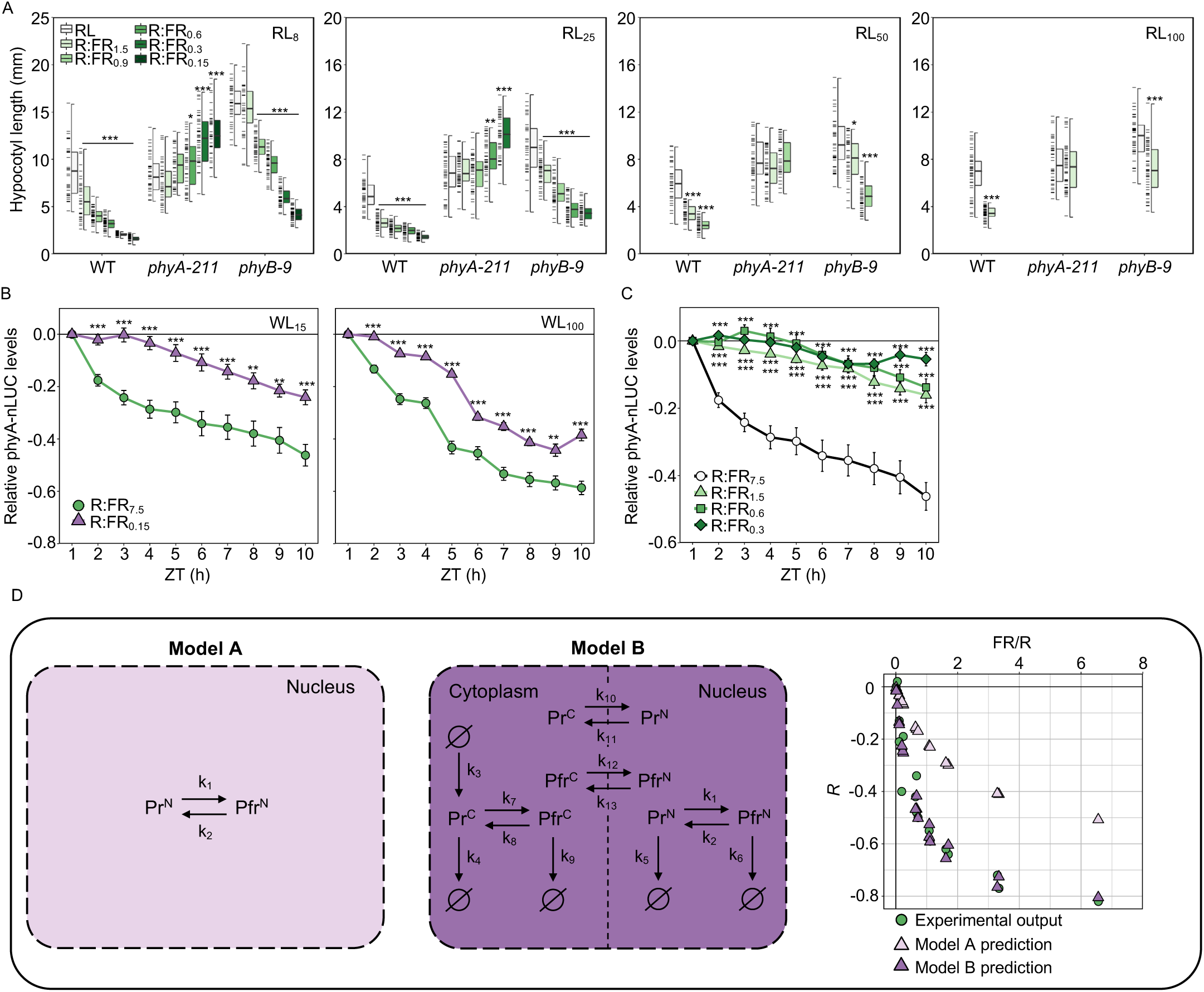
PhyA can respond to subtle changes in the R:FR ratio as predicted by modelling work. (*A*) Hypocotyl lengths of WT, *phyA-211* and *phyB-9* grown for 6 days in monochromatic red light (RL; 10L:14D) with or without supplementation of FR. Different RL intensities used (8, 25, 50 or 100 μmol m^-2^ s^-1^) are indicated in the top right corner of each panel. Supplemental FR was applied to each RL condition to reduce the R:FR to 1.5, 0.9, 0.6, 0.3 or 0.15. FR capabilities were limited at higher RL irradiances. Box plots show: median (centre line); upper and lower quartiles (box limits); 1.5 x interquartile range (whiskers); individual data points (dashes on the left of boxes, *n* ≥ 25). (*B*) Hourly relative phyA-nLUC abundance, calculated with respect to the first time point after dawn (ZT1) in high (R:FR_7.5_) or low (R:FR_0.15_) R:FR conditions applied at two WL intensities (PAR = 15 or 100 μmol m^-2^ s^-1^) and (*C*) under different R:FR ratios (R:FR = 7.5, 1.5, 0.6 or 0.3; PAR = 15 μmol m^-2^ s^-1^). Traces show mean signal produced by *n* ≥ 10 plants, error bars show mean ± SEM. Asterisks indicate significant differences from RL controls (*\*P* < 0.05; *\*\*P* < 0.01; *\*\*\*P* < 0.001), calculated using a Wilcoxon signed-rank test *(A*), and calculated using a Student’s t-test compared to R:FR_7.5_ controls (*B* and *C*). Each experiment was repeated three times. (*D*) Two alternative models (A, left-hand; B, right-hand) to explain phyA action. Pr^C^ and Pfr^C^ represent cytoplasmic phyA molecules in their respective states; Pr^N^ and Pfr^N^ show nuclear phyA molecules. Ø represents synthesis and degradation sources, and k_1-13_ reaction parameters. The plot to the right of the models shows the predicted (triangles) and actual (green circles, obtained from experimental work) relative hypocotyl length (*R*) values, plotted against a function of far-red to red light (FR/R).

We found that incremental reductions in R:FR gave rise to correlative increases in hypocotyl growth-repression in WT and *phyB-9*, but not in *phyA-211* (Fig. 2 *A*). This trend was observed across fluence rates, as was the promotion of phyA-nLUC by R:FR_0.15_ (Fig. 2 *A* and *B*). In contrast to phyA, which responds to a wide range of R:FR ratios, the phyB-driven elongation responses (evident in *phyA-211*) require more severe shade for induction (Fig. 2 *A*). Despite being described as a deep-shade detector (6, 8), we observed that a significant phyA-dependent growth inhibition occurred even at mild R:FR ratios, such as R:FR_1.5_ (Pfr/Ptot ∼0.71; Fig. 2 *A*, *SI Appendix*, Fig. S10 *A* and *B*, and Table. S1). In line with this, application of R:FR_1.5_ significantly increased phyA-nLUC abundance compared to control (R:FR_7.5_) conditions, although lower ratios of R:FR_0.6_ and R:FR_0.3_ did not lead to incremental rises in phyA-nLUC (Fig. 2 *C*). This highlights the importance of persistent subtle shade in boosting phyA levels and the phyA response. It also emphasises that the physiological response is not solely dictated by phyA levels but also involves other elements of the phyA signalling module (17).

Earlier studies have successfully applied modelling approaches as a tool to decipher the complex signalling properties of phyA (17, 19, 21). We aimed to determine if our experimental results align with the established theoretical frameworks describing phyA action. Specifically, we were interested in examining the relationship between irradiance and R:FR ratio as determinants of phyA function. To this end, we calculated the relative hypocotyl length of seedlings grown under descending R:FR (with respect to their lengths in the monochromatic RL), generated through variance in either FR or total (i.e. R+FR) intensity (raw data shown in Fig. 3 *A*). We established that there was a very weak correlation between FR (*R*^2^ 0.10, *p* = 0.17) and R (*R*^2^ 0.04, *p* = 0.26) fluence rate with hypocotyl growth inhibition, whereas a strong correlation exists with the R:FR (*R*^2^ 0.91, *p* < 0.001) (*SI Appendix*, Fig. S10 *B*).

**Fig. 3.**
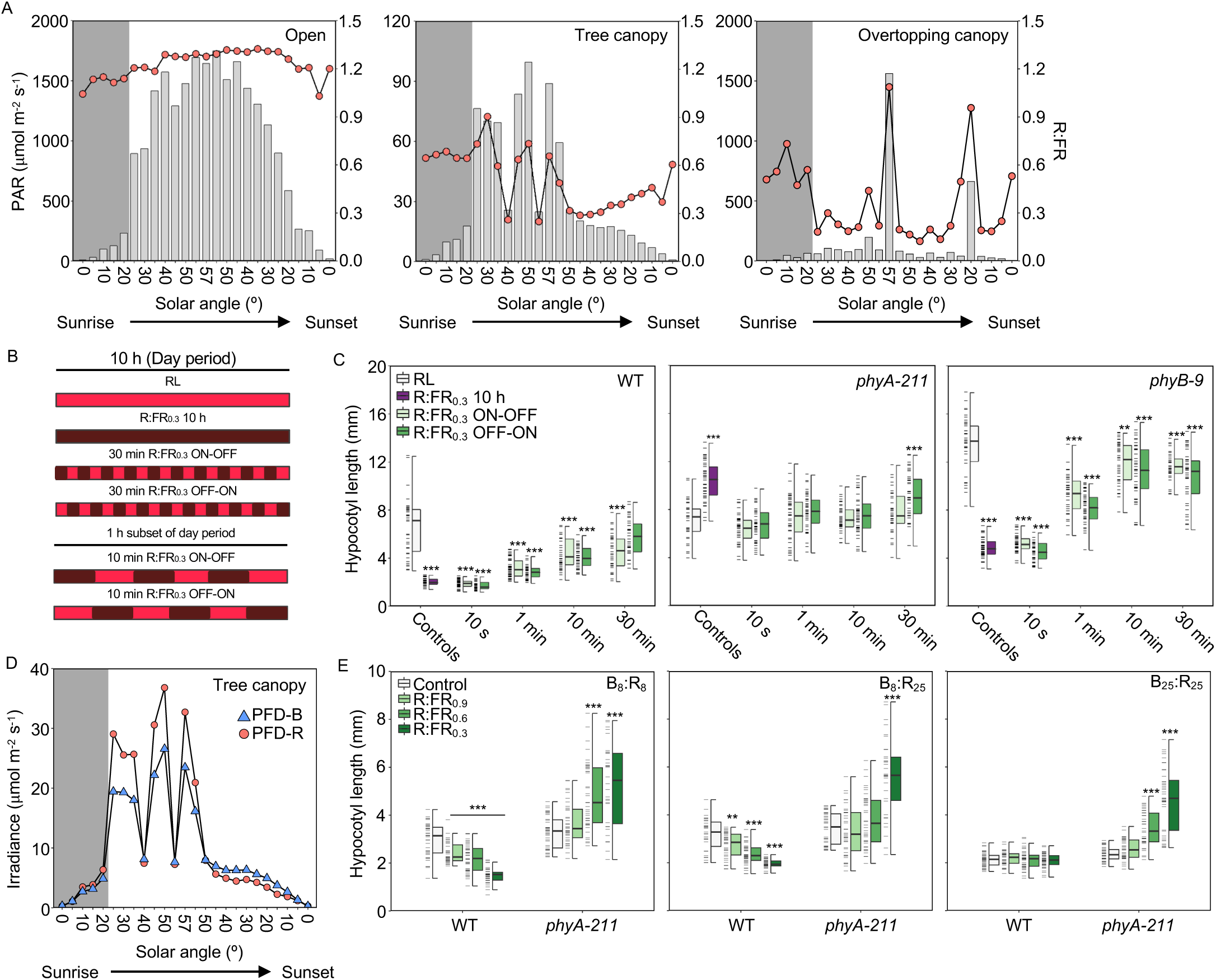
Spectral conditions within canopy shade are suited for phyA operation. (*A*) Comparison of PAR (light grey bars, left-hand y-axis) and R:FR ratio (600-700 nm:700-780 nm; red circles, right-hand y-axis) measurements across solar angle increments from sunrise to sunset in open (left), tree canopy shade (centre) and overtopping canopy (right) environments. (*B*) Visual representation of intermittent low R:FR treatments used in *C*. Each low R:FR period was followed by an equal length period without FR. ON-OFF treatments began the light period with low R:FR and finished without FR, while the opposite was true for OFF-ON. (*C*) Hypocotyl lengths of WT, *phyA-211* and *phyB-9* seedlings grown in monochromatic RL (25 μmol m^-2^ s^-1^) and either continuously (10 h), or intermittently (10 s, 1 min, 10 min or 30 min) treated with R:FR_0.3_ throughout the day period. (*D*) Photon flux density (PFD) of blue (B; 400-500 nm) and red (R; 600-700 nm) irradiances in canopy shade. Spectral data (*A* and *D*) were recorded on 4^th^ June 2022 near Leadburn, Scotland (UK; 55° 46’ 30.0” N, 3° 13’ 13.3” W). Dark grey background on each panel indicates ‘cloudy’ conditions. (*E*) Hypocotyl responses of WT and *phyA-211* mutants to a canopy shade relevant range of R:FR (0.3, 0.6 and 0.9) at combinations of 25 μmol m^-2^ s^-1^ B and R (B_25_:R_25_), reduced (8 μmol m^-2^ s^-1^) B irradiances (B_8_:R_25_) or reduced B and R (B_8_:R_8_). Box plots show: median (centre line); upper and lower quartiles (box limits); 1.5 x interquartile range (whiskers); individual data points (dashes on the left of boxes, *n* ≥ 25). Asterisks indicate significant differences from control conditions (*\*P* < 0.05; *\*\*P* < 0.01; *\*\*\*P* < 0.001). Significance was calculated using a Wilcoxon signed-rank test. Hypocotyl experiments were repeated three times.

To enable a direct comparison between model predictions and empirical hypocotyl data (*R*), we created an objective function, under the simple assumption that hypocotyl length linearly increases with phyA-Pfr nuclear levels. Our approach was to develop a series of model variants to test the functionality of key components of the phyA signalling module. The simplest model (Model A; Fig. 2 *D*), describes the switching of nuclear phyA between its Pr and Pfr forms. Model B is a more complex model which incorporates key cellular elements of the phyA signalling module. Adapted from Rausenberger *et al*. (17), Model B includes synthesis and degradation reactions and transport between the nuclear and cytosolic compartments facilitated by the FHY1/FHL transporters (Fig. 2 *D*). When compared to the experimental data, Model A predictions follow a similar trend, but fail to match the amplitude of hypocotyl length response (Fig. 3 *B*). In contrast, Model B provides an excellent fit to the data (Fig. 3 *B*). This indicates that elements of the phyA module in Model B, fulfilled by the photocycle-coupled nuclear shuttling, synthesis and degradation terms, are required to reproduce the full sensitivity range of hypocotyl growth inhibition observed in the experimental data (Fig 2 *A* and *D*). Lastly, we sought to establish the most parsimonious model that could accurately match the data. We found that a simplified model (Model C; *SI Appendix*, Fig. S10 *C*), which consolidated parameters yet preserved those related to nuclear flux and degradation rates, also provides a good qualitative match to the data. Our model-based analysis illustrates that steady-state phyA nuclear dynamics, which arises from the HIR response mode, provides an effective R:FR ratio sensing mechanism.

### Framing phyA activity in the context of natural shade

In order to contextualise these findings around the conditions experienced by plants in nature, we analysed how the R:FR ratio and PAR vary across a day in unshaded ‘open’ conditions and beneath two types of canopy shade: one produced by naturally occurring tree cover and one by ‘overtopping canopy’ closer to ground-level (simulated using four ∼ 30 cm tall *Solanum lycopersicum* plants; images displayed in *SI Appendix*, Fig. S11). PAR ranged from approximately 7-1749 μmol m^-2^ s^-1^ in ‘open’, 1-99 μmol m^-2^ s^-1^ beneath the ‘tree canopy’ and 2-1561 μmol m^-2^ s^-1^ under the ‘overtopping canopy’ (Fig. 3 *A* and *SI Appendix*, Table S2). While PAR diminished with solar angle in all habitats, we found that R:FR gradually increased in canopy shade toward dusk under clear skies. This appears to be a consequence of diminished light-reflectance as irradiances decrease, as there is a more apparent reduction in FR vs R intensity at low solar angles (*SI Appendix*, Fig. S12 *A* and Table S2). R:FR ratio ranged between 1.03-1.32 (mean = 1.23) in the ‘open’ environment, 0.25-0.90 (mean = 0.51) in ‘tree canopy’ and 0.14-1.09 (mean = 0.37) in the ‘overtopping canopy’ (Fig. 3 *A* and *SI Appendix*, Table S2). The estimated effects of the recorded spectra on the phytochrome-photoequilibria, both in terms of the k_1_/k_2_ transition and Pfr/Ptot ratios, are reported in Fig. S12 *B* and *C* and Table S3 (*SI Appendix*). The observed fluctuations in R:FR in both tree and overtopping canopy shade reflect the natural heterogeneity of these mild canopy environments, which causes fluctuations in Pfr/Ptot throughout the day (*SI Appendix*, Fig. S12 *C*). This is in part due to cloud cover variation: R:FR was lower beneath canopy shade under clear skies compared to cloudy conditions (grey background in Fig. 3 *A*; *SI Appendix*, Table S2) where FR is preferentially absorbed by atmospheric water vapour (42). In addition, we found spikes in PAR and R:FR ratio to regularly occur, most notably at midday (57°) in ‘overtopping canopy’, which are likely caused by sunlight penetrating through breaks in overstory foliage (sunflecks) (43, 44).

Given that phyA is able to detect mild canopy shade (Fig. 2 *A*) but is also reported to rely on continuous FR irradiation to operate in a HIR (14), we decided to test the robustness of phyA signalling during shade interruptions. To do this, R:FR_0.3_, simulating mild shade, was either provided continuously throughout a 10 h day period, or intermittently at 10 s, 1, 10, or 30 min intervals. Two alternating regimes were implemented: one terminating with high R:FR (ON-OFF) and one with low R:FR (OFF-ON) (Fig. 3 *B*). We found 10 s pulses were as effective as continuous R:FR_0.3_ in promoting phyA-repression of hypocotyl growth (Fig. 3 *C*). 1 min R:FR_0.3_ pulses also elicited a strong phyA response, while longer intervals between treatments were less effective (Fig. 3 *C*). Our data indicates that phyA is an adept sensor of R:FR in rapidly fluctuating light conditions, typical of natural mild canopy environments. Interestingly, phyB-driven hypocotyl elongation was only evident in *phyA-211* when 30 min pulses terminated with R:FR_0.3_ or under constant R:FR_0.3_ (Fig. 3 *C*). This suggests that, unlike phyA, phyB is a poor detector of intermittent mild shade.

In open environments, and in canopy shade sunflecks (R:FR_0.6-1.1_), the irradiance of R was generally greater than that of blue (B) wavelengths (Fig. 3 *D* and *SI Appendix*, Fig. S12 *D*). In modest, uninterrupted shade (i.e. R:FR_0.3_) this difference diminished, with R and B fluence rates dropping to around 8 μmol m^-2^ s^-1^ (Fig. 3 *D* and *SI Appendix*, Fig. S12 *D* and Table S2). Studies have shown that the depletion of B light operating through the cryptochrome photoreceptors, cry1 and cry2, can potentiate the effects of the phyB SAS (45, 46). We therefore wanted to test the impact of low B on the phyA R:FR ratio response. We found that phyA-mediated growth repression in response to reductions in R:FR ratio was greater in B_8_ than B_25_ (Fig. 3 *E*). In contrast, the addition of B did not have a notable impact on low R:FR-induced elongation in *phyA-211*. Thus, mild canopy shade conditions, characterised by low blue light (e.g. 8 μmol m^-2^ s^-1^) and low R:FR shade, appear to generate favourable conditions to enhance phyA hypocotyl suppression.

### In mild canopy shade phyA controls resource management and flowering time

Having established that phyA is a sensitive detector of modest canopy shade in seedlings, we subsequently extended our analysis to adult plants. We found exposure to R:FR_0.3_ significantly lengthened petioles of leaves 3 and 5 in *phyA-211* but not WT (Fig. 4 *A*). *phyB-9* petioles, on the other hand, were markedly shorter in R:FR_0.3_. Leaf blade size of all genotypes was similar in R:FR_8.5_ (Fig. 4 *B*). However, while WT and *phyA-211* did not change much in R:FR_0.3_, *phyB-9* became significantly smaller. We additionally noted a significant change in leaf dry weight for WT (−36%) and *phyB-9* (−46%) plants treated with R:FR_0.3_, which was less pronounced in *phyA-211* (−17%; Fig. 4 *B*). Consequently, phyA may limit leaf blade biomass in canopy shade by decreasing leaf thickness; indeed, the change in leaf mass per area (LMA) from R:FR_8.5_ to R:FR_0.3_ was smallest in *phyA-211* (Fig. 4 *B*). Therefore, phyA appears to modify leaf architecture under mild canopy shade conditions by suppressing petiole elongation and reducing leaf biomass, while overall leaf area is preserved.

**Fig. 4.**
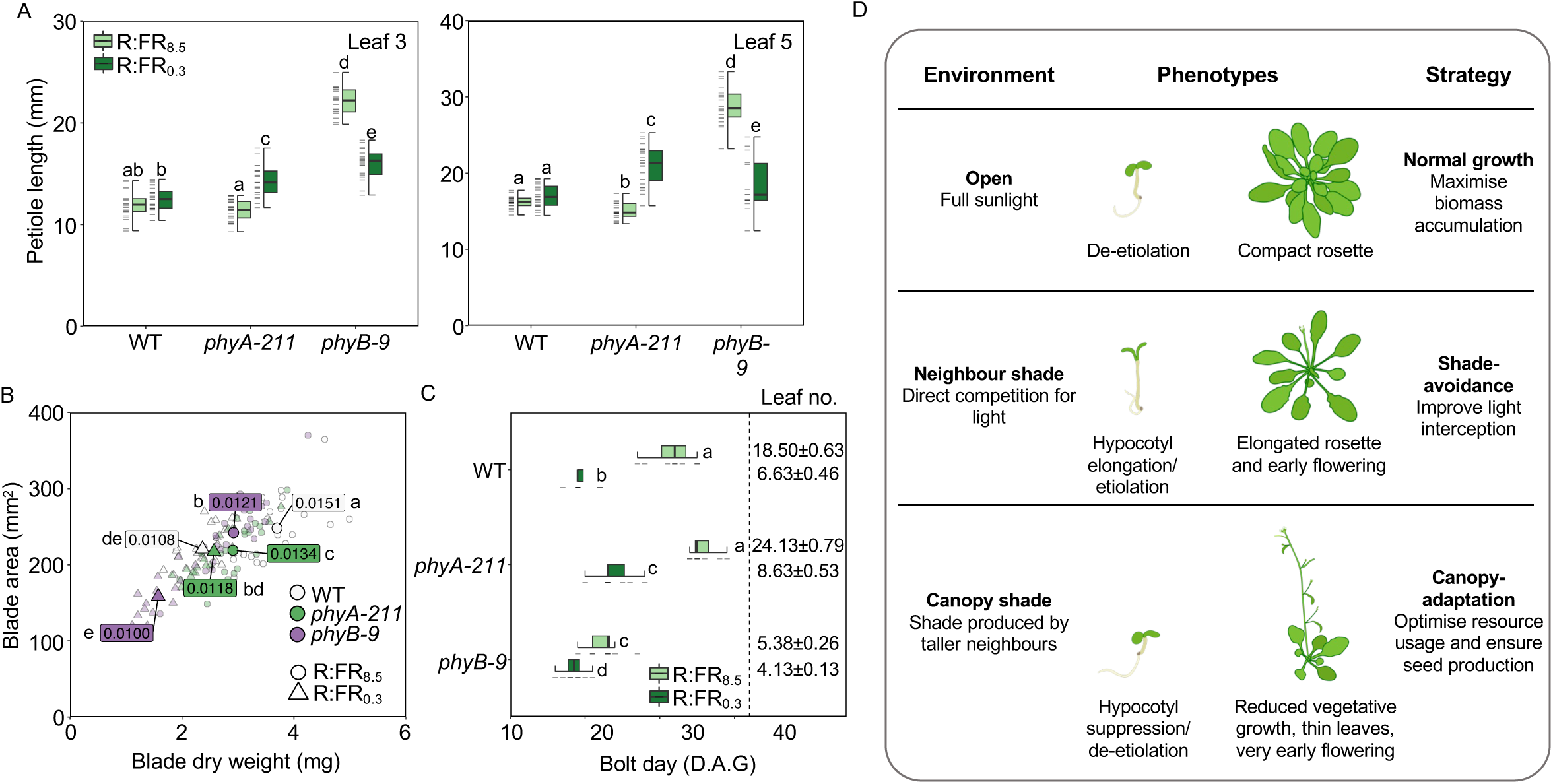
PhyA contributes to a distinct developmental strategy in adult plants that anticipates worsening shade. (*A*) Petiole length of leaf 3 and 5 from WT, *phyA-211* and *phyB-9* plants at harvest (35-days-old) following growth in WL (100 μmol m^-2^ s^-1^, 12L:12D) without (R:FR_8.5_; Pfr/Ptot ∼0.80) or with (R:FR_0.3_; Pfr/Ptot ∼0.35) FR supplementation. (*B*) Leaf area plotted against dry weight of leaf 5 at harvest. All data points are shown (transparent points), as well as the median for each genotype-treatment combination (solid points). Median leaf mass/area (LMA) is also labelled. Each treatment was repeated 3 times (*n* = 8), with the combined data from all repetitions being plotted (*n* ≥ 20 following outlier exclusion). (*C*) Median bolt day (box and whisker plots) and mean rosette leaf number of plants at bolting (± SEM; *n* = 8). Plants were grown in the same conditions as (*A*) and (*B*) but were not harvested after 35-days in (*C*). x-axis displays day after germination (D.A.G) at which bolting stem first appeared. Box plots show as follows: median (centre line); upper and lower quartiles (box limits); 1.5 x interquartile range (whiskers); individual data points (dashes on the left of boxes). Letters (*A-C*) denote statistically indistinguishable groups according to a Kruskal-Wallis test followed by a *post-hoc* Dunn’s Test (Bonferroni correction). (*D*) Summary of the growth strategies adopted by *Arabidopsis thaliana* in different shade and non-shade environments.

Accelerated flowering is a known important feature of phyB SAS, while both phyA and phyB operate in the regulation of photoperiodic flowering (12, 31, 47, 48). Consistent with this, R:FR_0.3_ accelerated flowering in WT, while the constitutively early flowering *phyB-9* showed a more subtle promotion by R:FR_0.3_ (Fig. 4 *C*). Conversely, the leaf number at bolting in R:FR_0.3_ was significantly higher in *phyA-211* compared to WT (Fig. 4 *C* and *SI Appendix*, Table S4). This data indicates that under mild shade conditions, flowering is hastened by both the deactivation of phyB and the action of phyA. This finding contrasts with the response of hypocotyl growth, where phyA acts to counteract the phyB SAS.

Our data collectively suggest that phyA has a broad and distinct role in plant adjustment to canopy shade, a phenomenon we have named the "Canopy Adaptation Strategy" (CAS). This strategy helps maintain seedling de-etiolation, conserve resources and accelerate the plant life cycle in environments where the adaptive value of the SAS is diminished (3, 5, 9), thereby improving survival chances (Fig. 4 *D*).

## Discussion

PhyA is a key photoreceptor regulating adaptive responses to natural shade. This research has shown that phyA is a sensitive detector of mild canopy shade and is able to reliably detect R:FR ratio even in heterogeneous conditions. PhyA appears to serve an important function in eliciting the canopy adaptation strategy (CAS), which optimises development in canopy shade conditions by preventing detrimental elongation growth habits in environments when shade cannot be outgrown.

Our study illustrates the highly dynamical properties of phyA which are strongly influenced by light and canopy shade conditions. It is well established that phyA is light labile, and depletes following light induced proteolysis (17, 19, 28). PhyA-nLUC analysis illustrates that proteasomal degradation is, to some extent, counterbalanced *PHYA* synthesis during a 10 h photoperiod (Fig. 1 *A* and *E*). This provides an explanation for the long-standing observation that basal levels of phyA are maintained through the light period (25; *SI Appendix*, Fig. S2). The persistence of phyA through the day may underlie responses such as photoperiodic flowering, particularly in plants grown in unshaded conditions (47, 48).

A key characteristic, that emerged from our analysis is that *PHYA* synthesis is incredibly responsive to persistent low R:FR. For instance, unlike the phyB low R:FR response, which is circadian gated (35), phyA levels and activity can be induced by prolonged exposure to low R:FR ratio at any time of day (Fig. 1 *C* and *D*). Additionally, our findings demonstrate that even slight decreases in the R:FR ratio, which are too subtle to drive the phyB SAS (Fig. 2 *A*; 4, 27), can increase phyA levels and activity. Therefore, beyond its established role as a deep shade sensor, the phyA sensory module appears to be tuned to detect mild canopy shade. We suggest this capability might enable plants to adapt their growth to shade conditions that may intensify over the course of a growing season. The capacity to anticipate increasing shade confers an adaptive advantage to plants, allowing them to complete their lifecycle and set seed before light availability becomes too limiting for growth.

Our data also uncovered modes of crosstalk between phyB and phyA. We discovered that the loss of phyB greatly enhances the activation of *PHYA* expression in response to shade light (Fig. 1 *H*). This sheds light on why phyA-induced hypocotyl inhibition is more pronounced in *phyB* mutants compared to the WT (Fig. 1 *C*; 8, 49). Thus, phyB appears to play an important role in moderating phyA growth suppression by preventing excessive *PHYA* accumulation in shade. Control of *PHYA* synthesis may be mediated through phyB suppression of PIFs, which are known to directly regulate *PHYA* transcription (32), or other factors such as those involved in chromatin modification (50). Similarly, the observed accelerated nighttime accumulation of phyA-nLUC in the *phyB* mutant (Fig. 1 *I* and *SI Appendix*, Fig. S9) may also arise, at least in part, from increased activity of PHYTOCHROME INTERACTING FACTOR4 (PIF4) and PIF5 which promote nocturnal *PHYA* expression through direct binding to the phyA promoter (32, 51).

Interestingly, *phyB-9* also alters the diel regulation of phyA protein abundance. Most notably, we observed enhanced depletion of phyA-nLUC during the daytime in *phyB-9* (Fig. 1 *I*), which can be inhibited by the proteasome inhibitor MG132 (Fig. 1 *G*). This suggests a role for phyB in decreasing the light lability of phyA through modulation of the proteasomal degradation pathways. Additional research is needed to confirm whether this mechanism is direct or involves post-transcriptional modifications, such as phosphorylation, which is known to modify phyA activity and its susceptibility to proteolysis (52). We hypothesise that connectivity between phyB and phyA likely plays a vital ecological role in varying light environments. For instance, the ability of phyB to exerts a moderating effect on *PHYA* expression may help dampen phyA responses in canopy shade. Equally, phyB enhances phyA protein stability in unshaded conditions, which may allow phyA to act in concert with phyB to promote photomorphogenic responses such as seedling de-etiolation.

PhyB is known to be an effective sensor of the R:FR ratio under full sunlight, where Pfr/Ptot is largely determined by the light spectrum and phyB is capable of detect changes in R:FR indicative of future shading by neighbours (22, 27). However, the reliability of phyB to act as a R:FR sensor diminishes in lower irradiance environments, such as those encountered beneath persistent canopy shade, where the role of phyB as a temperature sensor becomes more prominent (22). In contrast, the limited thermal reversion exhibited by phyA in many species, including various Arabidopsis accessions, may help facilitate accurate R:FR sensing across different climatic conditions (28, 29). However, while the phyA HIR is known to be fluence rate dependent, a property that is critical for detecting sustained low R:FR conditions experienced in canopy shade (Fig. 3 *A*), this could reduce the precision of phyA in detecting a range of R:FR ratios (8). We found that the extent of phyA-mediated hypocotyl growth inhibition only weakly correlated with irradiance, but strongly correlated with Pfr/Ptot calculated from R:FR ratio (Fig. 2 *A*, *SI Appendix*, Fig. S10 *A* and *B*, and Table S1). This suggests that the dependency on fluence rate enables the detection of canopy shade without significantly affecting the ability of phyA to sense the R:FR ratio. Modelling highlighted the relative importance of the phyA light reactions to this capability, including contributions from key elements of the HIR in the R:FR response (Fig. 3 *D*). Our simulations indicate that the light reactions are sufficient to trigger a response to shifts in the R:FR ratio (Model A, Fig 3. *D*). Linking the photocycle with the wider elements of the HIR is, however, essential to accurately reproduce the complete sensitivity spectrum to the R:FR ratio seen in the data (Model B, Fig. 3 *D*; 17).

Interestingly, we observed that low R:FR light boosts daytime phyA-nLUC primarily via increased synthesis rather than stabilisation of phyA (Fig. 1 *E* and *F*). Lowering the R:FR ratio is known to increase the proportion of phyA-Pr, the more stable phyA isomer, yet we observed a slight increase in phyA proteolysis (Fig. 1 *E* and *F*; 17, 28). This increase in proteolysis under low R:FR is likely due to an enhancement in the portion of the phyA pool within the nucleus, where it is targeted for degradation (25, 53). A previous modelling study highlighted the significance of phyA degradation in sustaining the phyA HIR response (17). As such, the combination of increased PHYA degradation, coupled with an upregulation of *PHYA* synthesis, may serve to enhance the flux and potentially amplify the phyA response under low R:FR ratios. Furthermore, it is noteworthy that while reducing R:FR to 1.5 stimulated a rise in phyA-nLUC, further reductions to 0.6 and 0.3 did not lead to successive rises in levels (Fig. 2 *C*). This again emphasises the importance of the phyA HIR module elements in delivering the R:FR ratio response (Fig. 2 *A* and *D*). We speculate that elevations in phyA may be important in eliciting a response to very mild shade where nuclear phyA would otherwise be low. Our modelling work (Fig. 2 *D*) also suggests that the nuclear concentrating HIR mechanism enables phyA to serve as a sensitive detector of R:FR ratio (*SI Appendix*, Fig. S10 *B*).

To contextualise the ecological relevance of our findings, we performed spectral analysis of two mild canopy shade environments across solar angles (*SI Appendix*, Table S3). R:FR ratio varied from 0.25-0.9 (estimated Pfr/Ptot ∼0.44-0.64) beneath the tree canopy and 0.12-0.19 (Pfr/Ptot ∼0.32-0.66) in the overtopping canopy environments (Fig. 3 *A* and *SI Appendix*, Tables S2 and S3), ranges within which we found phyA growth suppression to occur (Fig. 2 *A* and 3 *E*, *SI Appendix* Fig. S10 *B*). Our dataset highlighted the inherent heterogeneity of mild shade environments, which are regularly disrupted by sunflecks (43, 44, 54). When replicating such conditions in our lab experiments, we found that the phyA suppression response is maintained well with brief interruptions in mild shade (e.g. < 10 min), whereas the phyB SAS is not induced in intermittent mild shade (Fig. 3 *C*). Our results are consistent with previous modelling predictions indicating that phyB activity is dampened by quick transitions between shade and sunflecks, limiting its capability to detect R:FR shifts in canopy environments (22). This is believed to result from the cellular response speed, which may not keep pace with the rapidly changing light conditions. In contrast, phyA is better suited for functioning in heterogeneous light conditions, making it an effective detector of canopy shade.

Our study illustrates that phyA possesses a key role in regulating the response to canopy shade throughout the plant life cycle. Earlier research led to the proposal that the limited seed reserves in species like Arabidopsis are insufficient for shade-avoidance responses to occur, instead resulting in a preference for de-etiolation (55). In support of this, we found that even under very mild shade phyA inhibits hypocotyl elongation and maintains de-etiolation (Fig. 1 *C*, 2 *A* and 3 *E*). This highlights that de-etiolation is likely the default state when seedlings germinate in canopy shade. PhyA-driven de-etiolation in response to R:FR reductions are further facilitated by low blue light irradiances typically found in canopy shade (Fig. 3 *D* and *E*, and *SI Appendix*, Fig. S12 *D*). The adaptive value of this phyA-mediated response has been demonstrated previously, with *phyA* mutants having lower survival rates than WT in dense canopy shade (6). In adult plants, simulation of mild canopy shade from day 7 onwards promotes phyA-suppression of petiole elongation and leaf blade biomass reductions, while helping to maintain blade area, with this phenotype appearing to be dominant over phyB SAS (Fig. 4 *A* and *B*). These measures may be important for resource conservation and photosynthetic light-capture optimisation in irradiance limited, canopy shade environments (5, 9, 56). On the other hand, phyA works in concert with the phyB SAS to accelerate flowering in persistent canopy shade (Fig. 4 *C*).

Together, these results support previous studies which suggested that shade-induced elongation growth, such as the SAS, is maladaptive in rosette annuals growing beneath canopy shade (9, 57). We propose that phyA elicits a suite of responses in persistent shade, which we have named the Canopy Adaptation Strategy (CAS). This strategy enhances the survivability of seedlings by facilitating de-etiolation, restructuring carbon resource allocation, and shortening the plant lifecycle to boost the probability of seed-set (Fig. 4 *D*). In summary, the phyA photo-sensory system is precisely calibrated to sense canopy shade, activating the CAS to conserve resources and increase reproductive success.

## Methods

### Plant materials and growth conditions

Arabidopsis (*Arabidopsis thaliana*) *phyA-211* and *phyB-9* single, the *phyA-211 phyB-9* double were all of the Columbia-0 (Col-0) ecotype and have been described previously (58). The *PHYAp::LUC* (firefly LUCIFERASE) construct expressed in the Wassilewskija (Ws) genetic background (59).

Seeds were surface sterilised using liquid-phase surface sterilisation in bleach solution (20% [v/v] domestic bleach, 0.01% [v/v] Triton X-100) and stratified at 4°C in darkness for 72 h prior to sowing onto ½ Murashige and Skoog (MS) media (1.2% [w/v] agar, pH 5.7, no added sugars). For adult plant assays, seeds were sown directly onto F2+S Levington Advance Seed and Modular Compost plus sand soil mix (ICL Group, Israel) following stratification. In all experiments, 24 h prior to experiment start seeds received a 4 h white light (WL; 100 µmol m^-2^ s^-1^) pulse to synchronise germination, followed by 20 h darkness. Plants were grown at 22°C in all experiments. Seedling experiments were conducted using 10 h light:14 h dark (10L:14D) photoperiods, unless otherwise stated. Adult plant assays used a 12L:12D photoperiod and seedlings were grown using standard WL conditions (100 µmol m^-2^ s^-1^, R:FR = 8.5) and under a clear plastic lid for 6 days to ensure even germination. On day 7, the lid was removed, seedlings were thinned, and trays were transferred to the relevant condition.

WL was provided in Percival I30-BLL growth chambers (CFL Plant Climatics, Germany) by Luxline Plus F18W/840 fluorescent tubes (Sylvania, Newhaven, UK; spectrum shown in *SI Appendix*, Fig. S13 *A*). Neutral density filters (LEE Filters Worldwide, UK) were used to adjust WL irradiance to desired levels. Supplementary FR (peak ∼730 nm) was provided by OLSON 150 6+ Series FR LED strips (Intelligent LED Solutions, UK), filtered through LEE 120 Deep Blue colour filter (LEE Filters Worldwide) to remove excess R light. For experiments under monochromatic R ± FR and R + B ± FR, EB2-NE-PB Cooled Incubators (Snijders Labs, The Netherlands) custom fitted with ‘blue’ (peak ∼456 nm), ‘deep red’ (peak ∼680 nm) and ‘far-red’ (peak ∼782 nm) GreenPower LEDs (Philips, the Netherlands) were used (spectral outputs shown *SI Appendix*, Fig. S13 *B*). Light conditions were determined using a LI-180 Spectrometer (LI-COR, NE, USA). The total photosynthetic active radiation (PAR), R:FR ratios and the irradiances (photon flux density; PFD) of blue (B; 500-600 nm), green (G; 600-700 nm), red (R; 600-700 nm) and far-red (FR; 700-780 nm) wavebands used in experimental work are listed in Supplementary Table 5. R:FR was quantified as the ratio between wavelengths from 600-700 nm:700-780 nm. While narrower wavelength ranges are (e.g. 640-700 nm:700-760 nm and 640-670 nm:720-750 nm) are sometimes favoured as they lie closer to the absorption maxima of Pr (∼660 nm) and Pfr (∼730 nm) (12), we found that narrow waveband ranges resulted in lower R:FR calculations under broad-spectrum natural environments. This resulted in skewed Pfr/Ptot estimations when spectral data was integrated with phytochrome photoconversion cross-sections (60) compared to when R_600-700_:FR_700-780_ was used (*SI Appendix*, Table S3 and S6).

### Phytochrome photoequilibrium calculations

To estimate Pfr/Ptot, the spectral outputs of given light sources at set intervals were integrated with the Pr and Pfr conversion spectra of oat phyA (61). This returns the transition rate for Pr → Pfr (k_1_) and Pfr → Pr (k_2_) at any given wavelength and irradiation. Calculations were conducted using MATLAB (vR2024a) with scripts kindly provided by Dr Johanna Krahmer.

### Morphological data collection

Hypocotyl length data was collected from 6-day-old seedlings: images of seedlings were taken 1 h prior to dawn on their 7th treatment day and hypocotyl lengths were measured using ImageJ (http://rsbweb.nih.gov/ij/). Each treatment was repeated 3 times where *n* ≥ 25 for each genotype. Adult plants were harvested 1 h before dawn on day 36 of treatments (Fig. 5 *A* and *B*). Leaf 3 and 5 were removed, blades were severed from petioles and images were captured for respective measurements. All images were analysed using ImageJ. Blades of leaf 5 were wrapped in pre-weighed aluminium foil and dried at 80°C for 24 h before dry weight measurements were obtained. Leaf Mass per Area (LMA; mg/mm^2^) was calculated as leaf mass/leaf surface area. Time to flower (Fig. *5 C* and *SI Appendix,* Table S4) was taken as the day at which a bolting stem first became visible, at which point rosette leaf number was also recorded (Fig. 5 *C*).

### Spectral data collection

Spectral data from an un-shaded ‘open’ and woodland ‘canopy shade’ environment was collected on 4^th^ June 2022 near Leadburn, Scotland (UK; approximately 55° 46’ 30.0” N, 3° 13’ 13.3” W). This site was away from artificial street lighting and free of confounding obstructions to the path of light from the sun (i.e. from buildings and hills). Measurements were taken at 5° increments in solar angle (relative to the horizon) from sunrise to sunset (both 0°), as well as one recording at the solar zenith (57°). The corresponding time for each solar angle was determined for the nearest available location (Leadburn, UK) using an online resource (https://www.timeanddate.com/sun/). The presence/absence of cloud cover at each measurement point was noted: ‘cloudy’ conditions occurred when clouds visibly blocked the path of sunlight; other time points are marked ‘clear’. Spectral readings, measured as photon flux density (PFD; recorded in µmol m^-2^ s^-1^) across 1 nm increments from 380-780 nm, were collected using a LI-180 spectrometer (LI-COR, NE, USA). In addition, PAR as well as PFD-B (400-500 nm), -G (500-600 nm), -R (600-700 nm) and -FR (700-780 nm) were also obtained and used to calculate photon ratio measurements. Each measurement is an average of 6 readings taken across 1 min at 10 s intervals to account for small fluctuations in spectral composition.

### RNA extraction and cDNA synthesis

Seedlings were grown for 6 days in 10L:14D (PAR = 15 µmol m^-2^ s^-1^, R:FR = 7.5; 22°C). At the start of their 7^th^ day, seedlings were exposed to the relevant light conditions (WL [R:FR = 7.5] or FR [R:FR = 0.15]) where they were maintained until the relevant time points (ZT1, 3, 8 or 10), at which point seedling tissue was collected, snap frozen in liquid N_2_ and stored at −80°C. Frozen tissue was ground, then 300 µl of pre-heated (60°C) RE buffer (0.1 M Tris pH 8.0, 5 mM EDTA pH 8.0, 0.1 M NaCl, 0.5% SDS) + 1% β-mercaptoethanol was added and samples were vortexed until homogenised. 300 µl of a 1:1 acidic phenol:chloroform mix were added to the sample and vortexed further, before being transferred to ice. Samples were subsequently centrifuged at 4°C for 15 min at maximum speed. Up to 300 µl of the supernatant was transferred to a new collection tube, mixed gently, then stored at 4°C to induce the precipitation of nucleic acids. After 15 min, samples were centrifugated at 4°C for 30 min at maximum speed to pellet all nucleic acid precipitate. Supernatants were then discarded and pellets were gently washed with 300 µl of 70% ethanol (w/v), followed by a further centrifugation for 5 min at max speed (4°C). All traces of ethanol were removed and the pellet was dissolved in 50 µl of nuclease-free water. cDNA synthesis was conducted on extracted RNA using the qScript cDNA synthesis kit (Quantabio, MA, USA) as per the manufacturer’s instructions. cDNA concentration was quantified using an ND-1000 NanoDrop (Thermo Fisher Scientific, MA, USA).

### qRT-PCR

SYBR Green qPCR master-mix (Thermo Fisher Scientific) was used following the manufacturer’s protocol. Relative transcript abundance of *PHYA*, normalised to *PROTEIN PHOSPHATASE 2A* (*PP2A*), was calculated using the ΔΔCt-method. Gene-specific oligonucleotides for *PHYA* (FW-GTTTGGGACTGAGGAAGATGTG; RV-CTTTTGGGGACTACTTGTTTGC) and reference gene *PP2A* (FW-TAACGTGGCCAAAATGATGC; RV-GTTCTCCACAACCGCTTGGT) were used to quantify transcript levels. For each data point, three technical and three biological replicates were performed.

### Plasmid construction and plant transformation

To generate the phyA-nLUC (phyAp::phyA-nanoLUCx3FLAGx10His) transgenic line, the phyA promoter (2141 bp upstream of the inferred initiation codon) was amplified with Q5 High-Fidelity DNA polymerase (New England Biolabs, MA, USA) from the Arabidopsis Col-0 genome. The coding region was obtained from mRNA extracted using the RNAeasy mini kit (Qiagen, Germany). cDNA was synthesised using the qScript kit (Quantabio, MA, USA). The nLUC tag (NLUCx3FLAGx10His; NL3F10H) was amplified from the pET28::NL3F10H construct (61). All parts were recombined in the pGWB601 vector (62) using the Gibson Assembly method (NEBuilder HiFi DNA Assembly master mix kit). Primers for plasmid construction are listed in *SI Appendix*, Table S7. After sequence verification, *Escherichia coli* (DH5α) cells were transformed via heat-shock, following the Cold Spring Harbor Laboratory Protocol 1.24 (2006), and selected on spectinomycin plates. Plasmids were purified using the Monarch Plasmid Miniprep kit (New England Biolabs). Each cloned intermediate nucleotide sequence was verified through Sanger sequencing (Edinburgh Genomics, UK). *Agrobacterium tumefaciens* (strain ABI1, resistant to kanamycin and chloramphenicol) were transformed using the freeze and thaw method (63). Plant transformation was carried out using the floral-dip method (64). Segregation analysis of lines expressing the phyA-nLUC construct in the *phyA-211* mutant and the *phyA-211 phyB-9* double mutant background was performed using ½ MS plates containing 10 μg/ml Bialaphos. Phenotypic complementation assays were subsequently performed on homozygous lines, with *phyA-211* correctly expressing phyA-nLUC displaying a wild-type phenotype (*SI Appendix*, Fig. S1) and *phyA-211 phyB-9* for a *phyB-9* mutant.

### *In planta* bioluminescence assay

Surface-sterilised seeds were individually sown into wells of white LUMITRAC 96-well plates (Greiner Bio-One, Austria) containing 200 µl of solid ½ MS media prior to stratification for 72 h at 4°C. Seeds received a germination pulse (as described above) before a 7-day entertainment period began using 15 µmol m^-2^ s^-1^ WL (10L:14D). After this, 50 µl of 2% (v/v) Nano-Glo® (Promega, WI, USA) or 100 µl of 50 mM Luciferin (Sigma-Aldrich, MI, USA) was applied per well for nLUC and LUC assays, respectively. Plates were sealed with TopSeal-A plus (PerkinElmer, MA, USA) and transferred to a Tristar^2^ plate reader (Berthold, Germany) for 5-13 days, where bioluminescence was measured every hour with an integration time of 1.5 s per well. In between readings, seedlings were maintained in 15 µmol m^-2^ s^-1^ WL (Luxline Plus F18W/840, Sylvania) with (R:FR_0.15_) or without (R:FR_7.5_) supplemental FR (OLSON 150 6+ Series LED; Intelligent LED Solutions).

### Cycloheximide and MG132 treatment

Seedlings were initially grown in a 96-well plate for 4 days, following the protocol described for the bioluminescence assay. On the 5th day, seedlings were treated with the Nano-Glo® substrate and subjected to plate reader measurements. At the 9-day-stage, just before the final time point of the night period (ZT23), seedlings were treated with 200 µM CHX (C1988; Sigma-Aldrich) and/or 50 µM MG132 (474787; Sigma-Aldrich), with DMSO used as control. Bioluminescence readings from treated seedlings were subsequently taken every hour during the following light period (ZT1-10), with an integration time of 1.5 s per well.

### Model development

#### Model A

This consists of two reactions and two species (Pr^N^ and Pfr^N^). The rate at which Pr^N^ changes to Pfr^N^ is equal to 5.31 Nr + 0.82 Nfr while the rate at which Pfr^N^ changes to Pr^N^ is equal to 0.04 Nr + 1.70 Nfr where Nr and Nfr are the intensities of red and far-red light, respectively. The four constants in these expressions were extracted from (60) assuming the dominant wavelength of red light is 666 nm and that of far-red light is 730 nm. The standard deterministic rate equations (65) for this model were solved for the concentration of Pfr^N^ (denoted as [Pfr^N^]) as a function of the light intensities. Assuming the hypocotyl length is proportional to [Pfr^N^], it then follows that the quantity *R* = (L – L0)/L0 in Fig *3 B* of the main text is computed as ([Pfr^N^]- [Pfr^N^]*)/ [Pfr^N^]* where [Pfr^N^]* is the concentration of Pfr^N^ when the far-red light intensity Nfr is set to 0. Specifically, *R* is found to be given by equation (1).

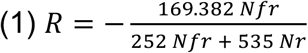

#### Model B

This consists of 13 reactions and four species (Pr^C^, Pfr^C^, Pr^N^ and Pfr^N^). The rates of the two reactions that change Pr to Pfr and vice versa (in cytoplasm or nucleus) after red or far-red absorption are as in Model A. We also set the transport rates from nucleus to cytoplasm (and vice versa) to be the same for Pr and Pfr. The decay rates of Pr and Pfr are also the same in the nucleus as in the cytoplasm. These assumptions considerably reduce the size of parameter space and hence simplify the optimization of the model to the data. The best fitting of the model’s prediction of *R* to the experimental data was implemented using the following procedure. We solve the rate equations for the model to obtain the concentration of Pfr^N^ and hence calculate *R* = (L – L0)/L0. The rate parameters which provide the closest fit were obtained by minimising the sum of the square of the differences between the experimental and predicted values of *R* where the sum is taken over the 22 measured tuples of Nfr, Nr and *R* values.

#### Model C (*SI Appendix*, Fig S10 *C*)

This consists of 6 reactions and two species (Pr^N^ and Pfr^N^). The rates of the two reactions that change one species to another after R or FR absorption are as in Model A. There are now four other rates (two for the production reactions and two for the removal reactions) which are unknown. We solve the rate equations for this model to obtain the concentration of Pfr^N^ and hence calculate *R* = (L – L0)/L0. The rate parameters which provide the closest fit were obtained by minimising the sum of the square of the differences between the experimental and predicted values of *R* where the sum is taken over the 22 measured tuples of Nfr, Nr and *R* values. Note that for simplicity and ease of finding the optimal parameter values, we reduced the size of parameter space by assuming that the rate of production of Pr^N^ is the same as that of Pfr^N^.

### Computational resources and data analysis

nLUC and LUC time-course data were processed through amplitude and baseline detrending, followed by −1,1 normalisation, using the BioDare software (www.biodare2.ed.ac.uk). Primers were designed utilising the ‘Primer Wizard’ tool available on the Benchling platform (www.benchling.com). Data were plotted using Rstudio (v.2024.04.1+748) and GraphPad Prism (v8.0) for Windows. Statistical methods are included in the figure legends. The model parameters that best fit the data were determined using Mathematica, Version 14.1 (Wolfram Research, Inc., Champaign, IL).

## Acknowledgements

This research was supported by an EastBio DTP BBSRC studentship awarded to PB, University of Edinburgh, School of Biological Sciences and CONACyT (CVU 537957) scholarships to MVC, and the Leverhulme Trust grant RPG-2024-082 awarded to KJH and RG. We thank Professor Millar’s lab (University of Edinburgh) and Dr Uriel Urquiza-Garcia (University of Düsseldorf) for providing the pET28::NL3F10H construct and NanoLUC protocols; and Dr Johanna Krahmer (University of Copenhagen) for the MATLAB scripts for calculating Pfr/Ptot at different R:FR ratios.

## Author contributions

PB, MVC, and KJH designed the experiments. PB and MVC conducted the experiments and analysed the data, with input from KJH. RG contributed by performing the modelling section. PB, MVC, and KJH drafted the manuscript, with additional input from RG.

## Notes

### Competing Interest Statement

The authors have declared no competing interest.

