## Supplementary Material for "*Arabidopsis thaliana* Phytochrome A Sensory Properties in Canopy Shade"

**SI appendix:**

Supplementary Figures
Supplementary Tables

### Supplementary Figures

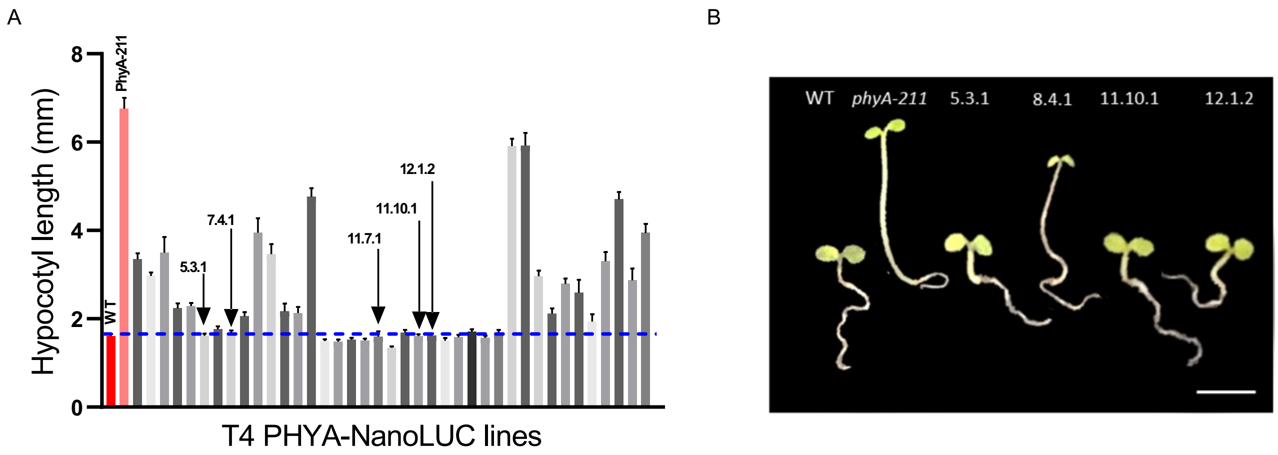


**Fig. S1. T4 selection of phyA-nanoLUC lines for phenotypic complementation of the *phyA-211* phenotype.** (*A*) Hypocotyl analysis of 36 phyA-nanoLUC homozygous lines (selected for resistance to bialaphos). Arrows indicate the first lines selected for analysis of PHYA protein dynamics. The blue line represents the length of WT (Col-0). (*B*) Representative image of 7-day-old seedlings as analysed on (*A*); scale bar = 1 cm. Seedlings were grown on ½ MS solid media at 22°C in 15 μmol $m^{-2}$ $s^{-1}$ (10L:14D) of white light supplemented with FR (R:FR = 0.15).

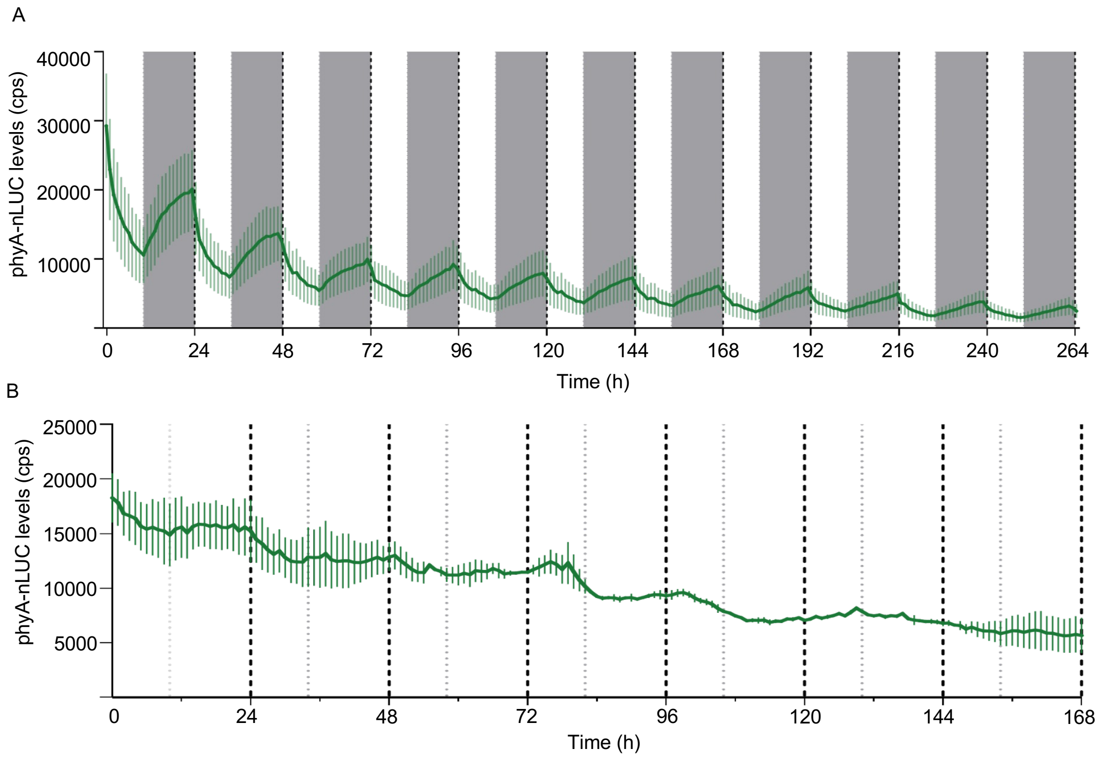


**Fig. S2. Bioluminescence analysis of phyA-nLUC under diurnal conditions (*A*) and constant light (*B*).** Dynamics of entrained phyA-nLUC seedlings grown in R:FR$_{7.5}$. Background colours of each panel correspond to R:FR$_{7.5}$ (white) and night (grey) periods. Bioluminescence was measured at 1 h intervals. Traces show mean signal produced by $n \geq 10$ plants; error bars show ± SEM.

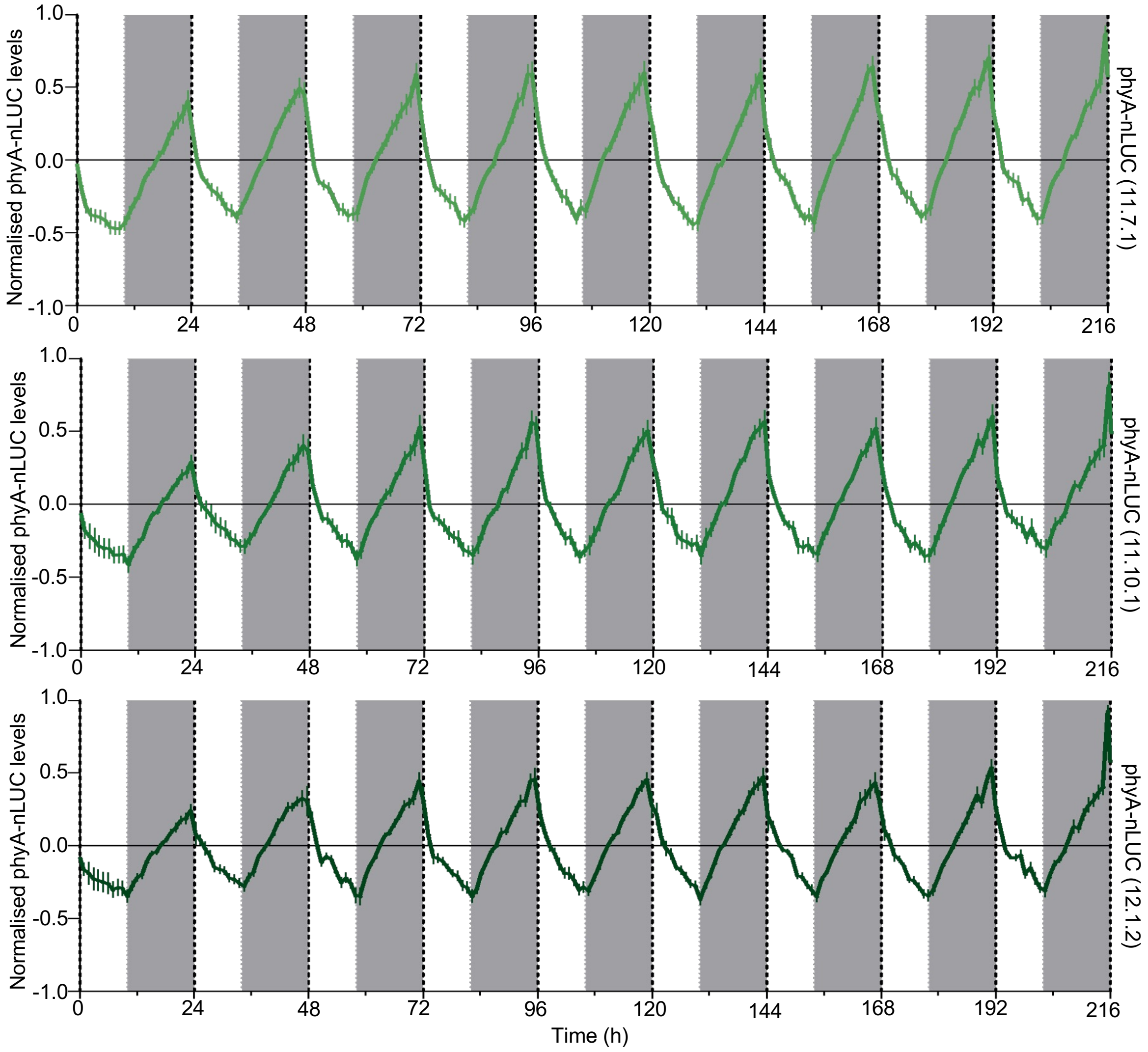


**Fig. S3. PhyA-nLUC diurnal dynamics tested on three different homozygous lines.** Normalised bioluminescence of phyA-nLUC of entrained seedlings grown in R:FR$_{7.5}$. Background colours of each panel correspond to R:FR$_{7.5}$ (white) and night (grey) periods. Lines are labelled on the right side of each panel. Bioluminescence was measured at 1 h intervals. Traces show mean signal produced by *n* ≥ 10 plants; error bars show ± SEM.

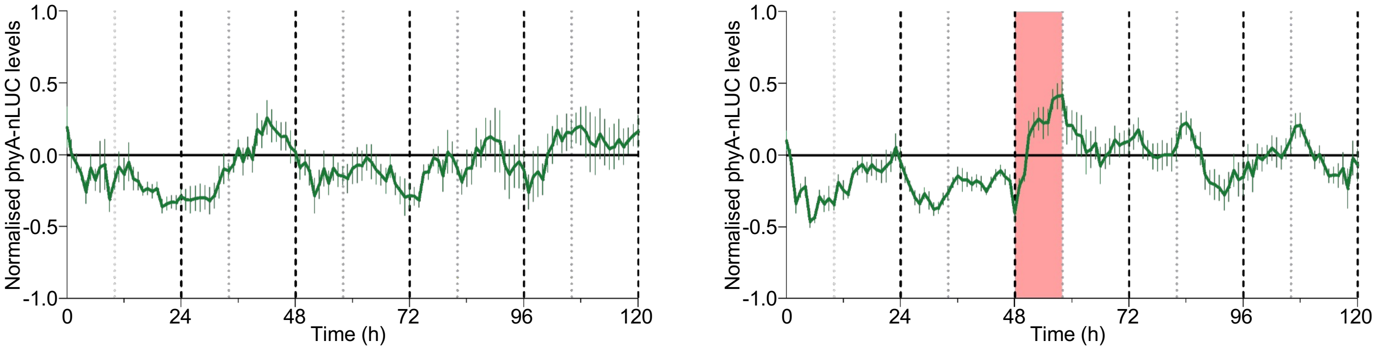


**Fig. S4 Low R:FR treatment increases phyA-nLUC levels in constant light.** Normalised bioluminescence of phyA-nLUC of entrained seedlings grown in R:FR$_{7.5}$ and transfer to constant light. Left-hand panel shows rhythms in constant R:FR$_{7.5}$, and right-hand panel shows R:FR$_{7.5}$ plus exposure to low R:FR (0.15) at 48 h. Background colours of each panel correspond to R:FR$_{7.5}$ (white) and R:FR$_{0.15}$ (red) periods. Bioluminescence was measured at 1 h intervals. Traces show mean signal produced by $n \geq 10$ plants; error bars show ± SEM.

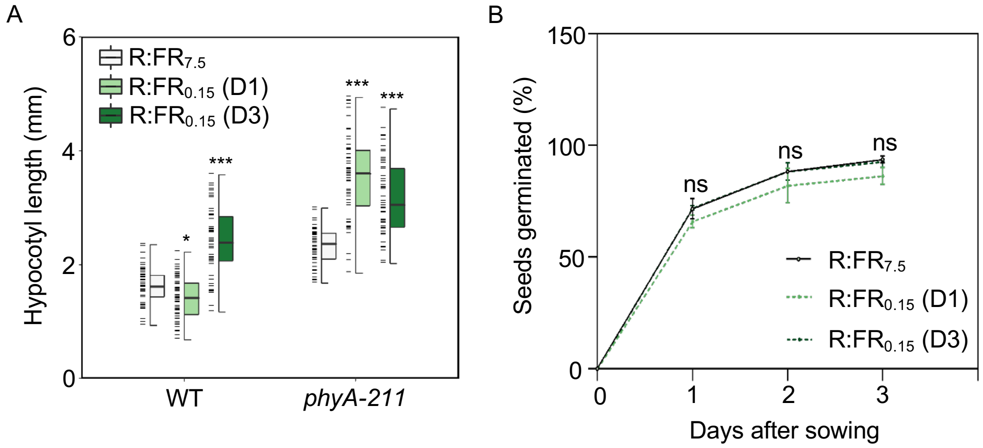


**Fig. S5 Hypocotyl and germination responses to application of low R:FR from day 1 or day 3.** (*A*) Hypocotyl lengths of 6-day-old WT and *phyA-211* seedlings grown in R:FR$_{7.5}$ (predicted Pfr/Ptot ~0.80) or R:FR$_{0.15}$ (~0.24) commencing either on day 1 (D1) or D3 post-germination pulse. PAR = 15 µmol m$^{-2}$ s$^{-1}$; 10L:14D. Germination pulses were initiated on 24 h prior to treatment commencement and consisted of 4 h 100 µmol m$^{-2}$ s$^{-1}$ WL (R:FR = 7.5) followed by 20 h darkness (22ºC). (*B*) Germination rates of WT and *phyA-211* seedlings in R:FR$_{7.5}$ (solid black line), R:FR$_{0.15}$ provided from day 1 (D1; dotted green line) or D3 (dotted black line) post-germination pulse (beginning on D0). Data in both *A* and *B* were produced in the same experiment. (*A*) Box plots display the median (centre line); upper and lower quartiles (box limits); 1.5 x interquartile range (whiskers); individual data points (dashes on the left of boxes, n ≥ 25). Asterisks indicate significant differences from controls (*$P < 0.05$; **$P < 0.01$; ***$P < 0.001$) calculated using a Wilcoxon signed-rank test. (*B*) Points indicate the mean; error bars show the SEM ($n \geq 25$). Statistical analysis was made using the Tukey's Method (one-way ANOVA; $P > 0.05$, ns).

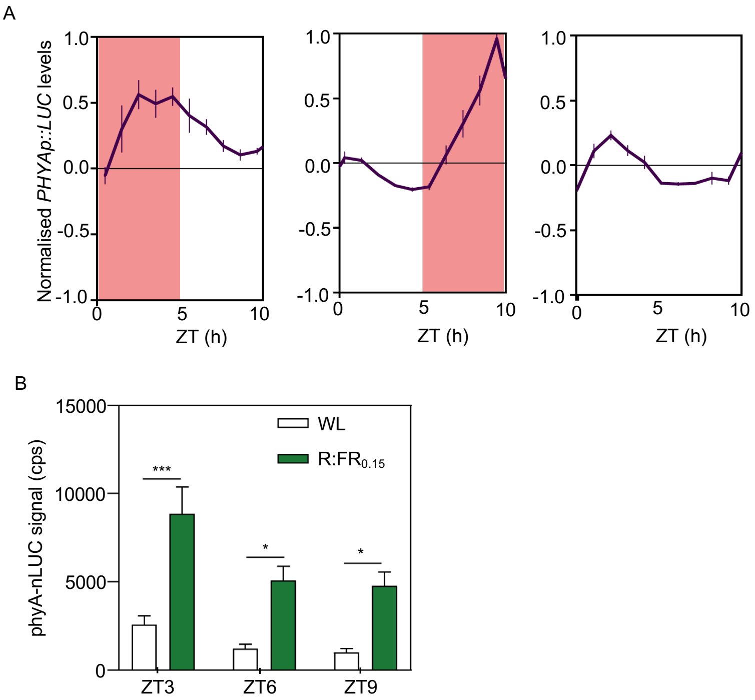


**Fig. S6. Low R:FR increases *PHYA* expression and phyA abundance throughout the day.** (*A*) Normalised bioluminescence of entrained *PHYAp::LUC* seedlings grown in R:FR$_{7.5}$ and exposed to low R:FR$_{0.15}$ in the morning (ZT0-5; left-hand panel), evening (ZT5-10; middle panel) or kept in R:FR$_{7.5}$ (right-hand panel). Background colours of each panel correspond to R:FR$_{7.5}$ (white) and R:FR$_{0.15}$ (red) periods. Bioluminescence was measured at 1 h intervals. Traces show mean signal produced by $n \geq 10$ plants; error bars show ± SEM. (*E*) phyA-nLUC signal strength at ZT3, 6 and 9 in R:FR$_{0.15}$ compared to the same time points in R:FR$_{7.5}$. Bars indicate the mean; error bars show ± SEM ($n \geq 10$). Asterisks indicate significant differences from controls (*$P < 0.05$; **$P < 0.01$; ***$P < 0.001$) calculated using a Student's t-test.

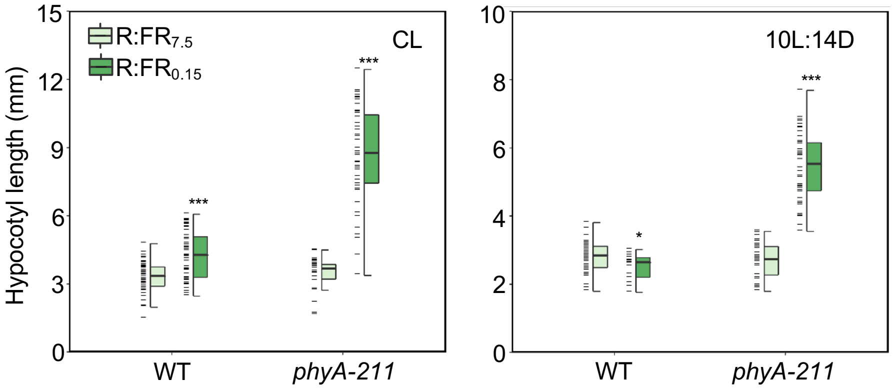


**Fig. S7. PhyA-mediated hypocotyl suppression is stronger in photoperiodic conditions.** Hypocotyl lengths of WT and *phyA-211* seedlings grown in R:$FR_{7.5}$, or R:$FR_{0.15}$ in constant light (CL; left-hand panel) or during a 10L:14D photoperiod (right-hand panel). PAR = 15 µmol $m^{-2}$ $s^{-1}$. Each box plot shows as follows: median (centre line); upper and lower quartiles (box limits); 1.5 x interquartile range (whiskers); individual data points (dashes on the left of boxes, $n \geq 25$). Asterisks indicate significant differences from controls (*$P$ < 0.05; **$P$ < 0.01; ***$P$ < 0.001) calculated using a Wilcoxon signed-rank test.

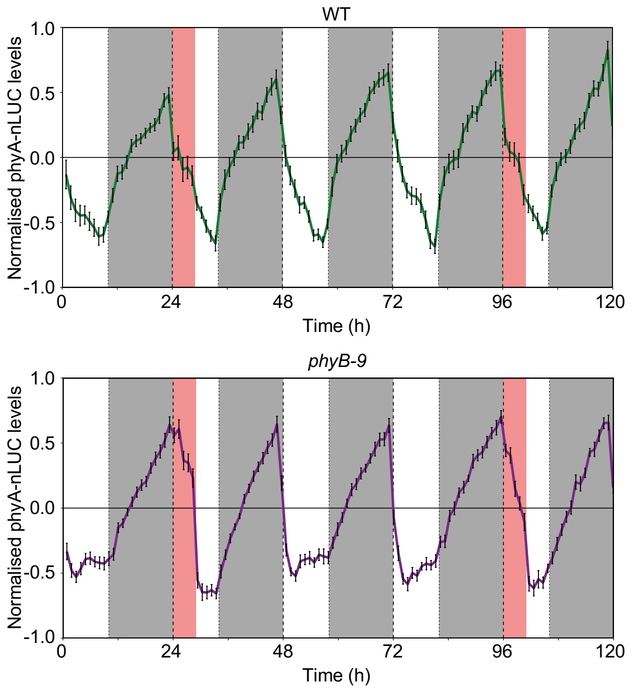


**Fig. S8. Change in phyA-nLUC levels during low R:FR periods is enhanced in a *phyB-9* background.** Normalised bioluminescence of phyA-nLUC across diurnal cycles in WT (green line, top panel) or a *phyB-9* (purple line, bottom panel) background. Two morning (ZT0-5) 5 h periods of R:$FR_{0.15}$ were provided: one from 24-29 h and one from 96-101 h. Traces show mean signal produced by $n \geq 12$ plants, measured at 1 h intervals; error bars show ± SEM. Background colours of each panel correspond to R:$FR_{7.5}$ (white), R:$FR_{0.15}$ (red) and night (grey) periods. All experiments were conducted in a background WL of 15 µmol $m^{-2}$ $s^{-1}$ (10L:14D). Experiments were repeated three times.

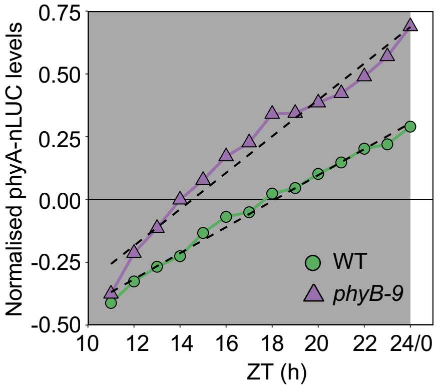


**Fig. S9. Loss of phyB accelerates night time accumulation of phyA-nLUC.** Normalised night-time bioluminescence of phyA-nLUC, produced in entrained WT (green circles) and *phyB-9* (purple triangles) seedlings between ZT10-24. Fitted regression lines for WT ($R^2$ = 0.99, $p$ < 0.001) and *phyB-9* backgrounds ($R^2$ = 0.97, $p$ < 0.001) had the following equations: WT, $y$ = 0.052($x$) - 0.939; *phyB-9*, $y$ = 0.073($x$) - 1.055. Traces represent mean values ± SEM ($n$ ≥ 10). Background colours of panels correspond to day (white) and night (grey) periods. Seedlings were entrained in 15 µmol $m^{-2}$ $s^{-1}$ WL ($R:FR_{7.5}$; 10L:14D) for 10 days prior to nLUC signal quantification. Experiments were repeated 3 times.

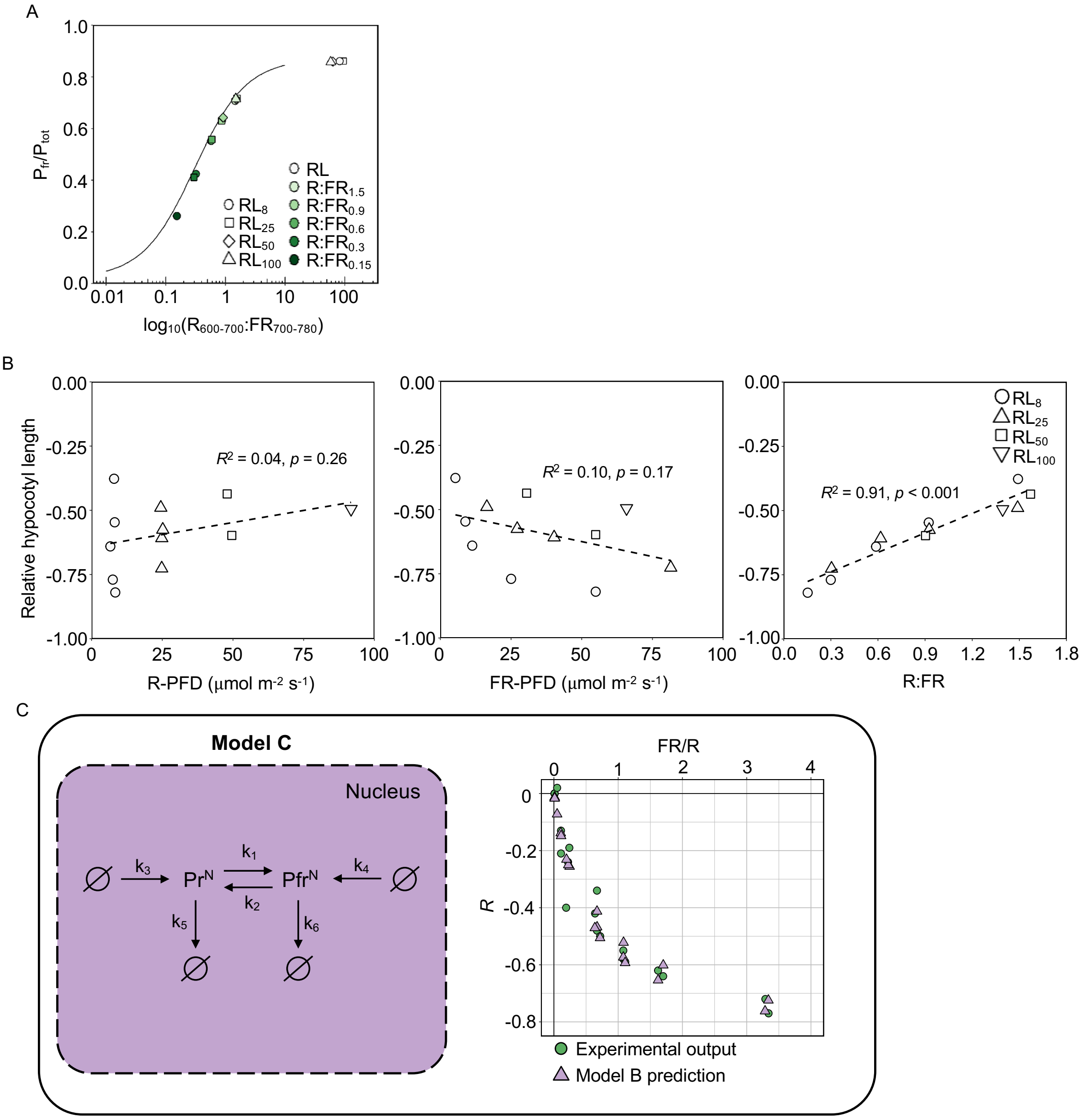


**Fig. S10. PhyA growth suppression correlates with external R:FR ratios.** (*A*) Predicted Pfr/Ptot ratios at corresponding R:FR (600-700 nm:700-780 nm) treatments used in experimental data shown in Fig. 3 *A*. R:FR ratios are plotted on a logarithmic scale. These predicted measurements correlate with the predicted phytochrome photoequilibrium from Mancinelli (1994), shown as the solid line, which was obtained using the R package photobiologyPlants (v0.4.1). (*B*) Median relative change in WT hypocotyl length (raw data shown in Fig. 3 *A*) under different R:FR ratios, calculated with respect to the average length of seedlings in the corresponding red light intensities. Linear regression analysis reveals a significant correlation between hypocotyl suppression and R:FR but not with R photon flux density (R-PFD) or FR-PFD. (*C*) Simplified version of Model B shown in Fig 3 *B*, which explains phyA growth suppression under R:FR conditions to a similar degree of accuracy. $Pr^N$ and $Pfr^N$ show nuclear phyA molecules. Ø represents synthesis and degradation sources, and $k_{1\text{-}6}$ reaction parameters. The plot to the right of the model shows the the predicted (triangles) and actual (green circles, obtained from experimental work) relative hypocotyl length (*R*) values, plotted against a function of far-red to red light (FR/R).

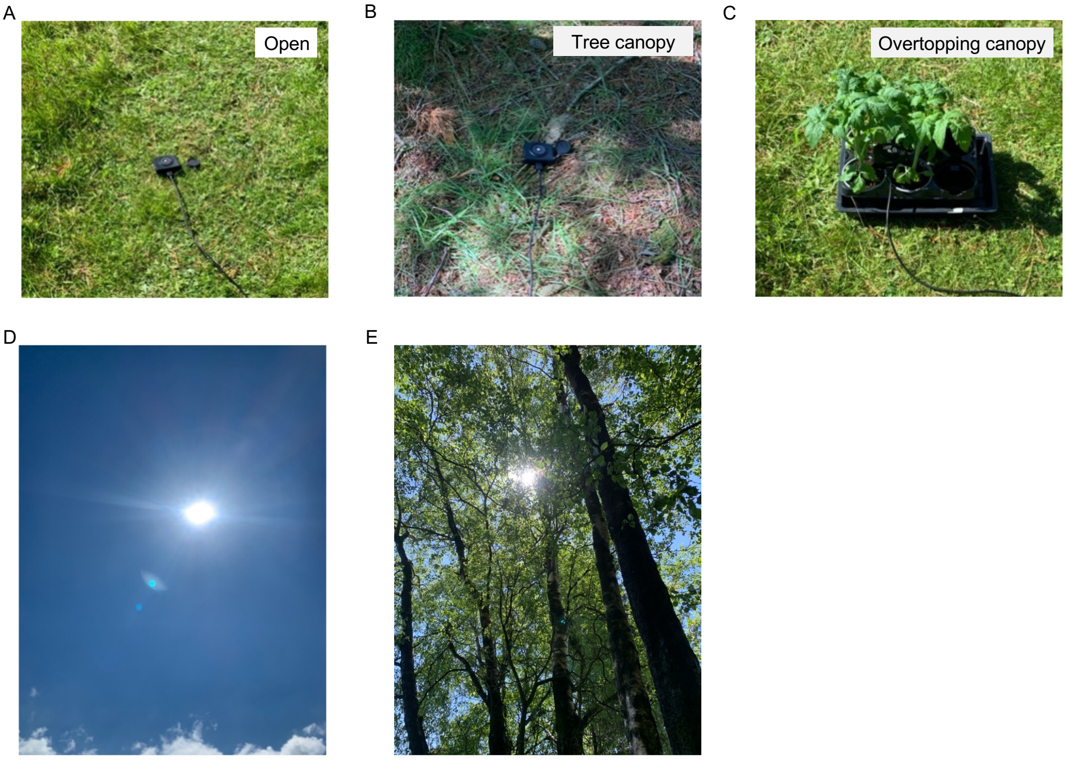


**Fig. S11. Images of the environments used for spectral data collection.** The open site had no overhead canopy cover (*A* and *D*) causing shading of the spectrometer at any point during the day. Two canopy shade environments consisted of a natural heterogenous woodland ('tree canopy'; *B* and *E*), which existed ~ 20 m from the 'open' measurement site or the addition of four ~ 30 cm tall *Solanum lycopersicum* plants to create an 'overtopping canopy' closer to ground-level (*C*). Images were taken ~ 13:00 on 4$^{th}$ June 2022, near Leadburn, Scotland (UK) (55° 46' 30.0" N, 3° 13' 13.3" W).

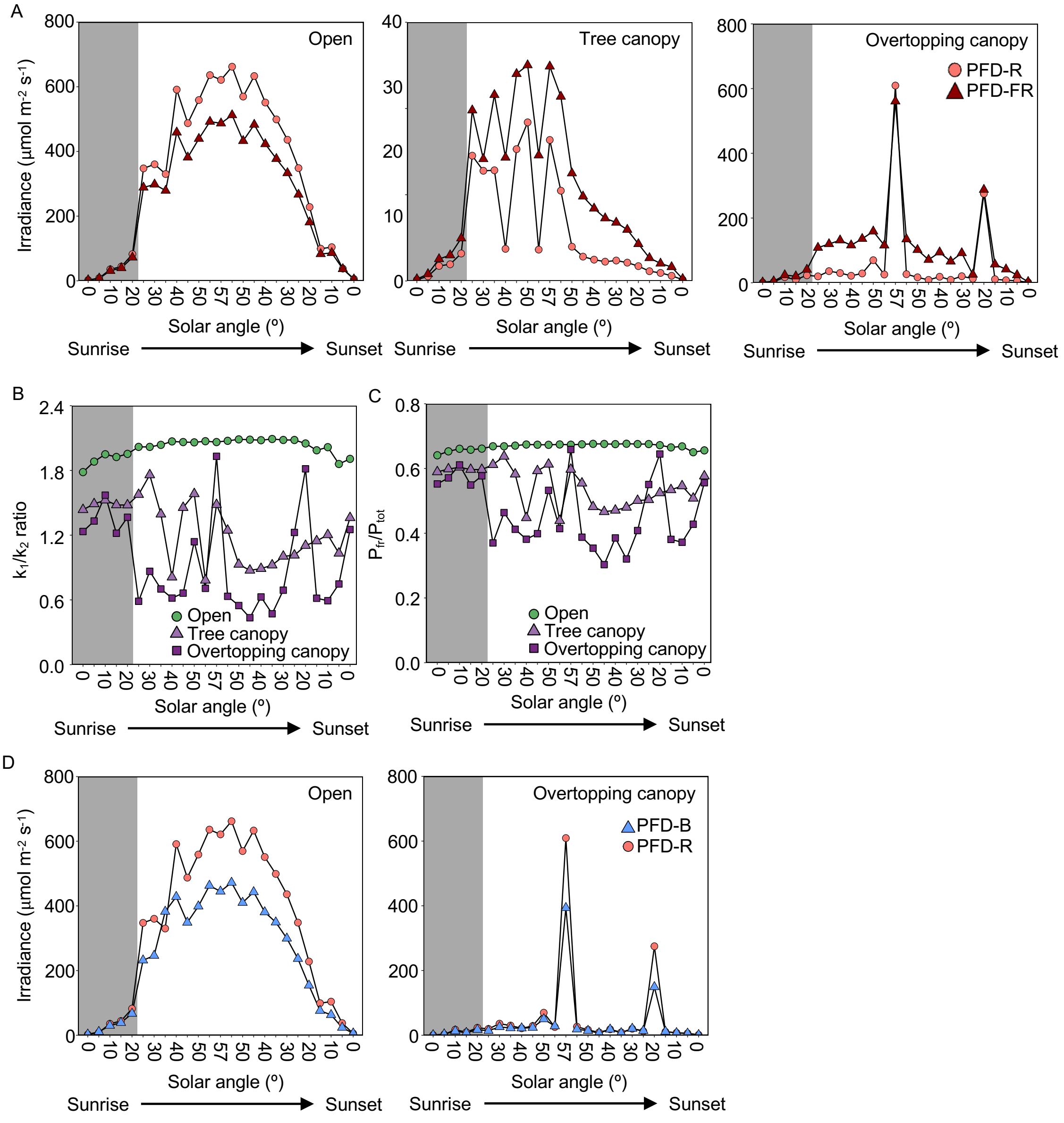


**Fig. S12. Variation in the spectral make up of open and canopy shade environments and their impacts on the phytochrome equilibrium**. (*A*) Comparison of the photon flux density (PFD)-red (R; 600-700 nm) vs PFD-far-red (FR; 700-780 nm) irradiances from sunrise to sunset in open and canopy shade environments. (*B*) Predicted ratios between $k_1$ (Pr → Pfr) and $k_2$ (Pfr → Pr) transition rate in natural open and canopy shade environments. Ratios were calculated using spectral irradiances (380-780 nm) integrated with the Pr and Pfr photoconversion spectra published in Mancinelli (1994). (*C*) Predicted Pfr/Ptot ratios from spectral data plotted against the $\log_{10}$ R:FR (600-700 nm:700-780 nm). Solid line indicates the predicted phytochrome photoequilibrium from Mancinelli (1994) calculated in monochromatic R and FR light, which was obtained using the R packaged photobiologyPlants (v0.4.1). (*D*) PFD-blue (B; 400-500 nm) vs PFD-R from the open and overtopping canopy environments; tree canopy shade data are plotted in Fig. 4 *E*. Dark grey panel background indicates cloudy weather conditions.

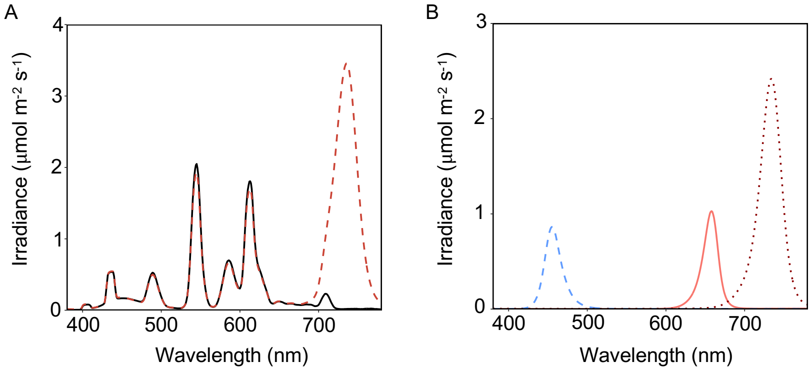


**Fig. S13. Spectral outputs of cabinets used in experimental work.** (*A*) Spectra of light in Percival I30-BL incubators fitted with Luxline Plus F18W/840 fluorescent tubes. Solid black line shows emission of fluorescent tubes only (i.e. WL or high R:FR); dashed red line shows the effects of additional FR supplementation using OLSON 150 6+ Series FR LED strips to generate a low R:FR (0.3) condition. (*B*) Spectral outputs of Phillips GreenPower 'blue' (dashed blue line), 'deep red' (solid red line) and 'far-red' (dotted dark red line) LEDs used to generate combinations of B ± R ± FR combinations.

### Supplementary Tables

**Table S1.** R:FR calculations across commonly used waveband definitions, along with corresponding phytochrome transition rate and Pfr/Ptot predictions for conditions used in **Fig. 2 *A*.**

**Table S2.** Summary of spectral data collected on 04.06.2022.

**Table S3.** R:FR calculations using different waveband definitions, along with corresponding phytochrome transition rate and Pfr/Ptot predictions from spectral data.

**Table S4.** Bolt time data from adult plant experiments.

**Table S5.** Spectral measurements from growth cabinets used in experimental work.

**Table S6.** R:FR calculations across commonly used waveband definitions from WL cabinets supplemented with FR LED lights, along with corresponding phytochrome transition rate and Pfr/Ptot predictions for data in Fig. 1 *C*, *H* and Fig. 4 *A-C*.

**Table S7.** Primers used for phyA-nLUC cloning.

**Table S1.** R:FR calculations across commonly used waveband definitions, along with corresponding phytochrome transition rate and Pfr/Ptot predictions for conditions used in **Fig. 2 *A***

| | | R:FR measurements | | | | |
|---|---|---|---|---|---|---|
| Red intensity (µmol $m^{-2}$ $s^{-1}$) | Treatment | 600-700nm: 700-780nm | 640-700nm: 700-760nm | 640-670nm: 720-750nm | $k_1/k_2$ transition ratio | Pfr/Ptot |
| 8 | | | | | | |
| | RL | 82.35 | 97.62 | 191.10 | 6.20 | **0.86** |
| | R:$FR_{1.5}$ | 1.46 | 1.32 | 1.71 | 2.41 | **0.71** |
| | R:$FR_{0.9}$ | 0.88 | 0.80 | 0.95 | 1.71 | **0.63** |
| | R:$FR_{0.6}$ | 0.58 | 0.53 | 0.59 | 1.23 | **0.55** |
| | R:$FR_{0.3}$ | 0.32 | 0.30 | 0.30 | 0.74 | **0.42** |

| | | | | | | |
|---|---|---|---|---|---|---|
| | R:FR$_{0.15}$ | 0.15 | 0.15 | 0.11 | 0.35 | **0.26** |
| 25 | | | | | | |
| | RL | 94.21 | 110.70 | 215.69 | 6.20 | **0.86** |
| | R:FR$_{1.5}$ | 1.56 | 1.41 | 1.72 | 2.53 | **0.72** |
| | R:FR$_{0.9}$ | 0.87 | 0.79 | 0.92 | 1.70 | **0.63** |
| | R:FR$_{0.6}$ | 0.59 | 0.54 | 0.60 | 1.26 | **0.56** |
| | R:FR$_{0.3}$ | 0.30 | 0.28 | 0.27 | 0.70 | **0.41** |
| 50 | | | | | | |
| | RL | 63.53 | 76.25 | 144.13 | 6.08 | **0.86** |
| | R:FR$_{1.5}$ | 1.51 | 1.38 | 1.62 | 2.50 | **0.71** |
| | R:FR$_{0.9}$ | 0.92 | 0.84 | 0.95 | 1.80 | **0.64** |
| 100 | | | | | | |
| | RL | 58.51 | 69.76 | 128.94 | 6.01 | **0.86** |
| | R:FR$_{1.5}$ | 1.50 | 1.39 | 1.55 | 2.50 | **0.71** |

**Note:** $k_1/k_2$ transition and Pfr/Ptot ratios were predicted according to Pr/Pfr conversion spectra of oat phyA, published by Mancinelli (1994). Calculations and scripts were obtained from Dr. Johanna Kramer.

**Table S2.** Summary of spectral data collected on 04.06.2022

| Environment | Solar angle | Approx. measurement time | Cloud cover | PFD measurements (µmols m⁻² s⁻¹) | | | | | | R:FR |
|---|---|---|---|---|---|---|---|---|---|---|
| | | | | PAR (400-700 nm) | UV (380-400nm) | Blue (400-500nm) | Green (500-600nm) | Red (600-700nm) | Far-red (700-780nm) | |
| Open | | | | | | | | | | |
| | 0° | 04:30 | Cloudy | 7.19 | 0.26 | 2.73 | 2.23 | 2.28 | 2.19 | 1.04 |
| | 5° | 05:21 | Cloudy | 28.40 | 1.07 | 10.70 | 9.50 | 8.39 | 7.41 | 1.13 |
| | 10° | 06:03 | Cloudy | 98.84 | 2.76 | 29.91 | 34.07 | 35.53 | 30.94 | 1.15 |
| | 15° | 06:43 | Cloudy | 125.76 | 3.70 | 38.61 | 44.13 | 43.90 | 39.42 | 1.11 |
| | 20° | 07:19 | Cloudy | 229.00 | 6.15 | 67.01 | 81.29 | 82.30 | 72.26 | 1.14 |
| | 25° | 07:54 | Clear | 892.77 | 18.90 | 231.94 | 319.51 | 347.57 | 288.51 | 1.20 |
| | 30° | 08:30 | Clear | 934.98 | 20.44 | 246.28 | 335.06 | 360.20 | 298.18 | 1.21 |
| | 35° | 09:06 | Clear | 1415.81 | 284.67 | 382.55 | 407.82 | 330.12 | 278.77 | 1.18 |
| | 40° | 09:44 | Clear | 1571.81 | 37.03 | 428.28 | 563.39 | 591.21 | 458.96 | 1.29 |
| | 45° | 10:24 | Clear | 1289.08 | 29.75 | 348.73 | 462.01 | 487.40 | 381.65 | 1.28 |
| | 50° | 11:07 | Clear | 1476.71 | 34.11 | 398.84 | 529.37 | 558.89 | 439.23 | 1.27 |
| | 55° | 12:06 | Clear | 1694.14 | 40.20 | 462.85 | 607.20 | 636.01 | 492.12 | 1.29 |
| | 57° | 13:11 | Clear | 1642.81 | 38.53 | 444.93 | 588.21 | 621.22 | 487.02 | 1.28 |
| | 55° | 14:00 | Clear | 1748.53 | 40.50 | 472.14 | 626.73 | 661.96 | 512.35 | 1.29 |
| | 50° | 15:06 | Clear | 1509.21 | 35.32 | 410.03 | 540.13 | 569.66 | 432.70 | 1.32 |
| | 45° | 15:55 | Clear | 1658.79 | 37.11 | 443.03 | 594.31 | 633.12 | 482.85 | 1.31 |
| | 40° | 16:33 | Clear | 1437.19 | 31.50 | 380.87 | 515.22 | 551.22 | 422.84 | 1.30 |
| | 35° | 17:09 | Clear | 1305.97 | 29.24 | 349.44 | 466.65 | 499.06 | 377.17 | 1.32 |
| | 30° | 17:47 | Clear | 1131.38 | 24.65 | 299.52 | 403.73 | 436.09 | 333.26 | 1.31 |
| | 25° | 18:21 | Clear | 898.01 | 19.42 | 236.66 | 319.24 | 348.41 | 266.73 | 1.31 |
| | 20° | 18:57 | Clear | 584.47 | 12.75 | 154.15 | 206.36 | 228.04 | 180.91 | 1.26 |

| | | | | | | | | | | |
|---|---|---|---|---|---|---|---|---|---|---|
| | 15° | 19:35 | Clear | 265.63 | 6.90 | 75.95 | 92.30 | 99.21 | 82.81 | 1.20 |
| | 10° | 20:12 | Clear | 251.79 | 4.97 | 63.03 | 86.52 | 103.95 | 86.16 | 1.21 |
| | 5° | 20:54 | Clear | 89.82 | 1.98 | 23.64 | 29.14 | 37.61 | 36.52 | 1.03 |
| | 0° | 21:41 | Clear | 16.86 | 0.53 | 6.01 | 5.36 | 5.60 | 4.67 | 1.20 |
| Tree canopy | | | | | | | | | | |
| | 0° | 04:30 | Cloudy | 0.88 | 0.03 | 0.32 | 0.29 | 0.28 | 0.43 | 0.65 |
| | 5° | 05:21 | Cloudy | 3.30 | 0.11 | 1.10 | 1.15 | 1.08 | 1.61 | 0.67 |
| | 10° | 06:03 | Cloudy | 9.59 | 0.25 | 2.73 | 3.44 | 3.49 | 5.09 | 0.69 |
| | 15° | 06:43 | Cloudy | 10.92 | 0.30 | 3.16 | 3.98 | 3.86 | 5.98 | 0.65 |
| | 20° | 07:19 | Cloudy | 17.60 | 0.45 | 4.87 | 6.48 | 6.37 | 9.90 | 0.64 |
| | 25° | 07:54 | Clear | 76.39 | 1.63 | 19.42 | 28.45 | 29.05 | 39.59 | 0.73 |
| | 30° | 08:30 | Clear | 70.12 | 1.70 | 19.30 | 25.76 | 25.56 | 28.28 | 0.90 |
| | 35° | 09:06 | Clear | 69.29 | 1.63 | 18.00 | 26.11 | 25.67 | 43.16 | 0.59 |
| | 40° | 09:44 | Clear | 25.62 | 0.99 | 8.11 | 10.24 | 7.45 | 28.63 | 0.26 |
| | 45° | 10:24 | Clear | 83.45 | 2.04 | 22.18 | 31.27 | 30.58 | 48.03 | 0.64 |
| | 50° | 11:07 | Clear | 99.47 | 2.37 | 26.57 | 36.80 | 36.79 | 50.05 | 0.74 |
| | 55° | 12:06 | Clear | 24.78 | 0.94 | 7.63 | 10.05 | 7.27 | 29.14 | 0.25 |
| | 57° | 13:11 | Clear | 88.90 | 2.15 | 23.48 | 33.34 | 32.70 | 49.76 | 0.66 |
| | 55° | 14:00 | Clear | 59.33 | 1.58 | 16.14 | 22.66 | 20.93 | 42.77 | 0.49 |
| | 50° | 15:06 | Clear | 25.91 | 0.96 | 7.99 | 10.15 | 7.94 | 24.98 | 0.32 |
| | 45° | 15:55 | Clear | 20.22 | 0.89 | 6.96 | 7.76 | 5.64 | 19.61 | 0.29 |
| | 40° | 16:33 | Clear | 17.91 | 0.80 | 6.28 | 6.82 | 4.94 | 16.82 | 0.29 |
| | 35° | 17:09 | Clear | 16.96 | 0.81 | 6.29 | 6.32 | 4.47 | 14.55 | 0.31 |
| | 30° | 17:47 | Clear | 17.38 | 0.78 | 6.33 | 6.43 | 4.74 | 13.55 | 0.35 |
| | 25° | 18:21 | Clear | 15.51 | 0.68 | 5.60 | 5.76 | 4.25 | 11.89 | 0.36 |
| | 20° | 18:57 | Clear | 13.07 | 0.58 | 4.96 | 4.76 | 3.44 | 8.59 | 0.40 |
| | 15° | 19:35 | Clear | 9.23 | 0.44 | 3.75 | 3.28 | 2.26 | 5.34 | 0.42 |

| | | | | | | | | | | |
|---|---|---|---|---|---|---|---|---|---|---|
| | 10° | 20:12 | Clear | 6.97 | 0.27 | 2.65 | 2.51 | 1.87 | 4.05 | 0.46 |
| | 5° | 20:54 | Clear | 3.85 | 0.12 | 1.32 | 1.35 | 1.20 | 3.24 | 0.37 |
| | 0° | 21:41 | Clear | 0.84 | 0.03 | 0.28 | 0.27 | 0.29 | 0.49 | 0.60 |
| Overtopping canopy | | | | | | | | | | |
| | 0° | 04:30 | Cloudy | 2.26 | 0.07 | 0.78 | 0.78 | 0.71 | 1.40 | 0.51 |
| | 5° | 05:21 | Cloudy | 9.15 | 0.34 | 3.29 | 3.29 | 2.63 | 4.71 | 0.56 |
| | 10° | 06:03 | Cloudy | 46.86 | 1.14 | 12.94 | 16.91 | 17.33 | 23.69 | 0.73 |
| | 15° | 06:43 | Cloudy | 27.39 | 0.72 | 7.56 | 10.48 | 9.54 | 20.13 | 0.47 |
| | 20° | 07:19 | Cloudy | 64.12 | 1.56 | 17.24 | 24.23 | 23.10 | 40.57 | 0.57 |
| | 25° | 07:54 | Clear | 60.56 | 1.55 | 13.99 | 27.25 | 19.72 | 108.41 | 0.18 |
| | 30° | 08:30 | Clear | 105.69 | 2.65 | 25.79 | 44.80 | 35.81 | 119.73 | 0.30 |
| | 35° | 09:06 | Clear | 94.54 | 2.58 | 22.37 | 42.82 | 29.99 | 131.25 | 0.23 |
| | 40° | 09:44 | Clear | 76.71 | 2.82 | 21.91 | 33.71 | 21.59 | 116.00 | 0.19 |
| | 45° | 10:24 | Clear | 95.81 | 2.87 | 23.53 | 44.17 | 28.75 | 135.86 | 0.21 |
| | 50° | 11:07 | Clear | 197.07 | 4.91 | 50.08 | 78.67 | 69.67 | 158.70 | 0.44 |
| | 55° | 12:06 | Clear | 92.63 | 3.61 | 28.19 | 39.56 | 25.51 | 115.62 | 0.22 |
| | 57° | 13:11 | Clear | 1560.71 | 31.50 | 393.69 | 568.70 | 609.32 | 560.61 | 1.09 |
| | 55° | 14:00 | Clear | 81.67 | 2.07 | 18.61 | 37.14 | 26.47 | 134.72 | 0.20 |
| | 50° | 15:06 | Clear | 57.89 | 1.80 | 14.11 | 27.38 | 16.78 | 101.66 | 0.17 |
| | 45° | 15:55 | Clear | 29.23 | 0.86 | 6.66 | 13.98 | 8.78 | 71.03 | 0.12 |
| | 40° | 16:33 | Clear | 66.68 | 2.45 | 18.18 | 30.44 | 18.49 | 94.15 | 0.20 |
| | 35° | 17:09 | Clear | 29.80 | 0.82 | 6.99 | 14.06 | 8.94 | 65.86 | 0.14 |
| | 30° | 17:47 | Clear | 71.42 | 2.55 | 20.59 | 31.11 | 20.20 | 91.81 | 0.22 |
| | 25° | 18:21 | Clear | 41.27 | 1.56 | 13.81 | 15.14 | 12.60 | 25.39 | 0.50 |
| | 20° | 18:57 | Clear | 662.26 | 9.99 | 149.24 | 242.64 | 275.02 | 287.51 | 0.96 |
| | 15° | 19:35 | Clear | 38.70 | 1.31 | 11.46 | 16.62 | 10.88 | 56.80 | 0.19 |

| | | | | | | | | | |
|---|---|---|---|---|---|---|---|---|---|
| 10° | 20:12 | Clear | 25.73 | 0.73 | 7.17 | 11.02 | 7.71 | 41.64 | 0.19 |
| 5° | 20:54 | Clear | 18.52 | 0.51 | 5.59 | 7.14 | 5.91 | 23.77 | 0.25 |
| 0° | 21:41 | Clear | 4.04 | 0.11 | 1.28 | 1.36 | 1.43 | 2.70 | 0.53 |

**Note:** R:FR was obtained by dividing PFD-R (600-700 nm) measurements by PFD-FR (700-780 nm).

**Table S3.** R:FR calculations across commonly used waveband definitions, along with corresponding phytochrome transition rate and Pfr/Ptot predictions from spectral data

| | | | | R:FR measurements | | | | |
|---|---|---|---|---|---|---|---|---|
| Environment | Solar angle | Approx. measurement time | Cloud cover | 600-700nm:700-780nm | 640-700nm:700-760nm | 640-670nm:720-750nm | k1/k2 transition ratio | Pfr/Ptot |
| Open | | | | | | | | |
| | 0° | 04:30 | Cloudy | 1.03 | 0.89 | 0.85 | 1.78 | **0.64** |
| | 5° | 05:21 | Cloudy | 1.12 | 0.91 | 0.90 | 1.88 | **0.65** |
| | 10° | 06:03 | Cloudy | 1.14 | 0.93 | 0.91 | 1.95 | **0.66** |
| | 15° | 06:43 | Cloudy | 1.10 | 0.89 | 0.87 | 1.92 | **0.66** |
| | 20° | 07:19 | Cloudy | 1.13 | 0.90 | 0.88 | 1.95 | **0.66** |
| | 25° | 07:54 | Clear | 1.19 | 0.95 | 0.94 | 2.02 | **0.67** |
| | 30° | 08:30 | Clear | 1.20 | 0.95 | 0.94 | 2.02 | **0.67** |
| | 35° | 09:06 | Clear | 1.22 | 0.97 | 0.95 | 2.04 | **0.67** |

| | | | | | | | | |
|---|---|---|---|---|---|---|---|---|
| | 40° | 09:44 | Clear | 1.28 | 1.00 | 1.00 | 2.07 | **0.67** |
| | 45° | 10:24 | Clear | 1.26 | 1.00 | 0.99 | 2.06 | **0.67** |
| | 50° | 11:07 | Clear | 1.26 | 0.99 | 0.99 | 2.06 | **0.67** |
| | 55° | 12:06 | Clear | 1.28 | 1.01 | 1.00 | 2.07 | **0.67** |
| | 57° | 13:11 | Clear | 1.26 | 1.00 | 0.99 | 2.06 | **0.67** |
| | 55° | 14:00 | Clear | 1.28 | 1.01 | 1.00 | 2.07 | **0.67** |
| | 50° | 15:06 | Clear | 1.30 | 1.02 | 1.02 | 2.09 | **0.68** |
| | 45° | 15:55 | Clear | 1.30 | 1.02 | 1.02 | 2.09 | **0.68** |
| | 40° | 16:33 | Clear | 1.29 | 1.02 | 1.01 | 2.08 | **0.68** |
| | 35° | 17:09 | Clear | 1.31 | 1.03 | 1.03 | 2.09 | **0.68** |
| | 30° | 17:47 | Clear | 1.30 | 1.02 | 1.01 | 2.08 | **0.68** |
| | 25° | 18:21 | Clear | 1.29 | 1.02 | 1.02 | 2.08 | **0.68** |
| | 20° | 18:57 | Clear | 1.25 | 1.00 | 0.98 | 2.05 | **0.67** |
| | 15° | 19:35 | Clear | 1.19 | 0.95 | 0.93 | 1.99 | **0.67** |
| | 10° | 20:12 | Clear | 1.19 | 0.98 | 0.97 | 2.02 | **0.67** |
| | 5° | 20:54 | Clear | 1.02 | 0.88 | 0.84 | 1.86 | **0.65** |
| | 0° | 21:41 | Clear | 1.19 | 1.00 | 0.97 | 1.91 | **0.66** |
| Tree canopy | | | | | | | | |
| | 0° | 04:30 | Cloudy | 0.64 | 0.57 | 0.51 | 1.44 | **0.59** |
| | 5° | 05:21 | Cloudy | 0.66 | 0.57 | 0.52 | 1.49 | **0.60** |
| | 10° | 06:03 | Cloudy | 0.68 | 0.58 | 0.53 | 1.52 | **0.60** |
| | 15° | 06:43 | Cloudy | 0.64 | 0.54 | 0.49 | 1.48 | **0.60** |
| | 20° | 07:19 | Cloudy | 0.64 | 0.54 | 0.49 | 1.48 | **0.60** |
| | 25° | 07:54 | Clear | 0.73 | 0.60 | 0.55 | 1.58 | **0.61** |
| | 30° | 08:30 | Clear | 0.89 | 0.73 | 0.68 | 1.76 | **0.64** |
| | 35° | 09:06 | Clear | 0.59 | 0.49 | 0.44 | 1.40 | **0.58** |
| | 40° | 09:44 | Clear | 0.26 | 0.21 | 0.17 | 0.81 | **0.45** |
| | 45° | 10:24 | Clear | 0.63 | 0.52 | 0.47 | 1.46 | **0.59** |

| | | | | | | | | |
|---|---|---|---|---|---|---|---|---|
| | 50° | 11:07 | Clear | 0.73 | 0.60 | 0.55 | 1.58 | **0.61** |
| | 55° | 12:06 | Clear | 0.25 | 0.20 | 0.16 | 0.78 | **0.44** |
| | 57° | 13:11 | Clear | 0.65 | 0.54 | 0.49 | 1.48 | **0.60** |
| | 55° | 14:00 | Clear | 0.48 | 0.41 | 0.35 | 1.24 | **0.55** |
| | 50° | 15:06 | Clear | 0.31 | 0.26 | 0.22 | 0.93 | **0.48** |
| | 45° | 15:55 | Clear | 0.28 | 0.23 | 0.19 | 0.87 | **0.47** |
| | 40° | 16:33 | Clear | 0.29 | 0.24 | 0.20 | 0.89 | **0.47** |
| | 35° | 17:09 | Clear | 0.30 | 0.25 | 0.21 | 0.92 | **0.48** |
| | 30° | 17:47 | Clear | 0.34 | 0.29 | 0.24 | 1.00 | **0.50** |
| | 25° | 18:21 | Clear | 0.35 | 0.29 | 0.25 | 1.01 | **0.50** |
| | 20° | 18:57 | Clear | 0.40 | 0.33 | 0.28 | 1.10 | **0.52** |
| | 15° | 19:35 | Clear | 0.42 | 0.35 | 0.30 | 1.15 | **0.53** |
| | 10° | 20:12 | Clear | 0.45 | 0.38 | 0.34 | 1.20 | **0.55** |
| | 5° | 20:54 | Clear | 0.37 | 0.33 | 0.28 | 1.03 | **0.51** |
| | 0° | 21:41 | Clear | 0.60 | 0.55 | 0.47 | 1.36 | **0.58** |
| Overtopping canopy | | | | | | | | |
| | 0° | 04:30 | Cloudy | 0.50 | 0.45 | 0.38 | 1.23 | **0.55** |
| | 5° | 05:21 | Cloudy | 0.55 | 0.47 | 0.41 | 1.33 | **0.57** |
| | 10° | 06:03 | Cloudy | 0.72 | 0.61 | 0.56 | 1.57 | **0.61** |
| | 15° | 06:43 | Cloudy | 0.47 | 0.40 | 0.34 | 1.22 | **0.55** |
| | 20° | 07:19 | Cloudy | 0.56 | 0.47 | 0.42 | 1.37 | **0.58** |
| | 25° | 07:54 | Clear | 0.18 | 0.15 | 0.11 | 0.59 | **0.37** |
| | 30° | 08:30 | Clear | 0.29 | 0.24 | 0.20 | 0.86 | **0.46** |
| | 35° | 09:06 | Clear | 0.22 | 0.18 | 0.14 | 0.70 | **0.41** |
| | 40° | 09:44 | Clear | 0.18 | 0.14 | 0.11 | 0.62 | **0.38** |
| | 45° | 10:24 | Clear | 0.21 | 0.16 | 0.13 | 0.66 | **0.40** |
| | 50° | 11:07 | Clear | 0.43 | 0.36 | 0.30 | 1.14 | **0.53** |

| | | | | | | | |
|---|---|---|---|---|---|---|---|
| 55° | 12:06 | Clear | 0.22 | 0.17 | 0.14 | 0.71 | **0.41** |
| 57° | 13:11 | Clear | 1.08 | 0.87 | 0.83 | 1.93 | **0.66** |
| 55° | 14:00 | Clear | 0.19 | 0.16 | 0.12 | 0.63 | **0.39** |
| 50° | 15:06 | Clear | 0.16 | 0.12 | 0.09 | 0.55 | **0.35** |
| 45° | 15:55 | Clear | 0.12 | 0.10 | 0.07 | 0.43 | **0.30** |
| 40° | 16:33 | Clear | 0.19 | 0.15 | 0.11 | 0.63 | **0.39** |
| 35° | 17:09 | Clear | 0.13 | 0.11 | 0.08 | 0.47 | **0.32** |
| 30° | 17:47 | Clear | 0.22 | 0.17 | 0.13 | 0.69 | **0.41** |
| 25° | 18:21 | Clear | 0.49 | 0.40 | 0.35 | 1.23 | **0.55** |
| 20° | 18:57 | Clear | 0.95 | 0.78 | 0.73 | 1.82 | **0.64** |
| 15° | 19:35 | Clear | 0.19 | 0.15 | 0.12 | 0.62 | **0.38** |
| 10° | 20:12 | Clear | 0.18 | 0.15 | 0.11 | 0.59 | **0.37** |
| 5° | 20:54 | Clear | 0.24 | 0.21 | 0.17 | 0.75 | **0.43** |
| 0° | 21:41 | Clear | 0.52 | 0.47 | 0.40 | 1.25 | **0.56** |

**Note:** k1/k2 transition and Pfr/Ptot ratios were predicted according to Pr/Pfr conversion spectra of oat phyA, published by Mancinelli (1994). Calculations and scripts were obtained from Dr. Johanna Kramer.

**Table S4.** Bolt time data from adult plant experiments

| Genotype | Treatment | Median bolt day | % plants flowered at harvest (D36) | Significance group |
|---|---|---|---|---|
| WT | | | | |
| | R:$FR_{8.5}$ | 34 | 45.8 | a |
| | R:$FR_{0.3}$ | 23 | 100 | b |
| *phyA-211* | | | | |
| | R:$FR_{8.5}$ | ND | ND | --- |
| | R:$FR_{0.3}$ | 27 | 100 | c |
| *phyB-9* | | | | |
| | R:$FR_{8.5}$ | 25 | 100 | d |
| | R:$FR_{0.3}$ | 20.5 | 100 | e |

**Note:** No *phyA-211* plants flowered by day 36 (D36). Significance between median bolt day calculated using *post-hoc* Dunn's Test with Bonferroni corrections.

**Table S5.** Spectral measurements from growth cabinets used in experimental work

| Corresponding figures | Background light | Treatment name | Irradiance (μmol m⁻² s⁻¹) PAR (400-700 nm) | B (400-500nm) | G (500-600nm) | R (600-700nm) | FR (700-780nm) | R:FR |
|---|---|---|---|---|---|---|---|---|
| **Fig. 1** | | | | | | | | |
| | $WL_{15}$ | | | | | | | |
| | | $WL/R:FR_{7.5}$ | 15.68 | 2.47 | 6.55 | 6.74 | 0.88 | 7.64 |
| | | $R:FR_{0.15}$ | 14.91 | 2.28 | 5.69 | 7.02 | 50.82 | 0.14 |
| **Fig. 4** | | | | | | | | |
| | $WL_{100}$ | | | | | | | |
| | | $R:FR_{8.5}$ | 100.83 | 21.53 | 43.24 | 36.53 | 4.3 | 8.51 |
| | | $R:FR_{0.3}$ | 97.94 | 20.9 | 40.95 | 36.56 | 119.35 | 0.31 |
| **Fig. 3 *A*** | | | | | | | | |
| | $RL_{8}$ | | | | | | | |
| | | RL | 8.12 | < 0.5 | < 0.5 | 8.04 | < 0.5 | 87.36 |
| | | $R:FR_{1.5}$ | 8.08 | < 0.5 | < 0.5 | 7.95 | 5.33 | 1.49 |
| | | $R:FR_{0.9}$ | 8.36 | < 0.5 | < 0.5 | 8.17 | 8.85 | 0.92 |
| | | $R:FR_{0.6}$ | 6.76 | < 0.5 | < 0.5 | 6.63 | 11.28 | 0.59 |
| | | $R:FR_{0.3}$ | 7.76 | < 0.5 | < 0.5 | 7.5 | 25.03 | 0.3 |
| | | $R:FR_{0.15}$ | 8.9 | < 0.5 | < 0.5 | 8.38 | 54.94 | 0.15 |
| | $RL_{25}$ | | | | | | | |
| | | RL | 26.24 | < 0.5 | < 0.5 | 26.02 | < 0.5 | 93.33 |
| | | $R:FR_{1.5}$ | 24.77 | < 0.5 | < 0.5 | 24.48 | 16.44 | 1.49 |
| | | $R:FR_{0.9}$ | 25.72 | < 0.5 | < 0.5 | 25.17 | 27.18 | 0.93 |

| | | | | | | | |
|---|---|---|---|---|---|---|---|
| | $R:FR_{0.6}$ | 25.27 | < 0.5 | < 0.5 | 24.83 | 40.24 | 0.62 |
| | $R:FR_{0.3}$ | 25.68 | < 0.5 | < 0.5 | 24.8 | 81.48 | 0.3 |
| $RL_{50}$ | | | | | | | |
| | RL | 50.47 | < 0.5 | < 0.5 | 50 | 0.55 | 90.45 |
| | $R:FR_{1.5}$ | 48.71 | < 0.5 | < 0.5 | 47.95 | 30.51 | 1.57 |
| | $R:FR_{0.9}$ | 50.32 | < 0.5 | < 0.5 | 49.59 | 54.93 | 0.9 |
| $RL_{100}$ | | | | | | | |
| | RL | 97.99 | < 0.5 | 0.68 | 97.11 | 1.04 | 93.27 |
| | $R:FR_{1.5}$ | 93.26 | 0.52 | 0.84 | 91.77 | 65.85 | 1.4 |
| **Fig. 3 *C*** | | | | | | | |
| $RL_{25}$ | | | | | | | |
| | RL | 24.27 | < 0.5 | < 0.5 | 23.92 | < 0.5 | 66.67 |
| | $R:FR_{0.3}$ | 26.50 | < 0.5 | < 0.5 | 25.71 | 78.94 | 0.33 |
| **Fig. 3 *E*** | | | | | | | |
| $B_8:R_8$ | | | | | | | |
| | Control | 15.96 | 8.21 | < 0.5 | 7.57 | < 0.5 | 49.06 |
| | $R:FR_{0.9}$ | 15.98 | 8.40 | < 0.5 | 7.40 | 8.55 | 0.87 |
| | $R:FR_{0.6}$ | 15.64 | 8.10 | < 0.5 | 7.34 | 12.70 | 0.58 |
| | $R:FR_{0.3}$ | 15.88 | 8.02 | < 0.5 | 7.62 | 26.67 | 0.29 |
| $B_8:R_{25}$ | | | | | | | |
| | Control | 32.52 | 7.66 | < 0.5 | 24.65 | < 0.5 | 85.36 |
| | $R:FR_{0.9}$ | 33.17 | 7.49 | < 0.5 | 25.4 | 28.76 | 0.88 |
| | $R:FR_{0.6}$ | 32.92 | 7.5 | < 0.5 | 25.12 | 41.02 | 0.61 |
| | $R:FR_{0.3}$ | 35.16 | 7.67 | < 0.5 | 27.08 | 88.5 | 0.31 |

| | | | | | | | |
|---|---|---|---|---|---|---|---|
| $B_{25}$:$R_{25}$ | | | | | | | |
| | Control | 50.86 | 25.42 | < 0.5 | 25.02 | < 0.5 | 97.01 |
| | R:$FR_{0.9}$ | 51.31 | 25.26 | 0.54 | 25.54 | 28.82 | 0.89 |
| | R:$FR_{0.6}$ | 52.79 | 25.44 | 0.59 | 26.78 | 44.52 | 0.6 |
| | R:$FR_{0.3}$ | 51.32 | 25.23 | 0.71 | 25.41 | 88 | 0.29 |

**Table S6.** R:FR calculations across commonly used waveband definitions from WL cabinets supplemented with FR LED lights, along with corresponding phytochrome transition rate and Pfr/Ptot predictions for data in Fig. 1 *C*, *H* and Fig. 4 *A*-*C*.

| | | R:FR measurements | | | | |
|---|---|---|---|---|---|---|
| PAR (μmols $m^{-2}$ $s^{-1}$) | Treatment | 600-700nm: 700-780nm | 640-700nm: 700-760nm | 640-670nm: 720-750nm | $k_1/k_2$ transition ratio | Pfr/Ptot |
| 15 | | | | | | |
| | R:$FR_{7.5}$ | 7.76 | 1.27 | 4.63 | 4.02 | **0.80** |
| | R:$FR_{0.15}$ | 0.15 | 0.04 | 0.02 | 0.32 | **0.24** |
| 100 | | | | | | |
| | R:$FR_{8.5}$ | 8.81 | 1.36 | 5.03 | 3.97 | **0.80** |
| | R:$FR_{0.3}$ | 0.30 | 0.06 | 0.04 | 0.55 | **0.35** |

**Note:** k1/k2 transition and Pfr/Ptot ratios were predicted according to Pr/Pfr conversion spectra of oat phyA, published by Mancinelli (1994). Calculations and scripts were obtained from Dr. Johanna Kramer.

**Table S7.** Primers used for phyA-nLUC cloning

| Gene part | Primer Sequence (5' - 3') |
|---|---|
| PHYAp FW | TCACTATGGCGGCCCCTCGATGCAACATGGGCCATGACTA |
| PHYAp RV | TCGGCCTAGAGCCTGACATTTTTTTTCCTGACACAGAGACA |
| PHYA CDS FW part 1 | GTCTCTGTGTCAGGAAAAAAAATGTCAGGCTCTAGGCCGA |
| PHYA CDS RV part 1 | ATCCATCTCATAGTCCTTCCAAGGTAAACTCCTTGTCTTGAC |
| PHYA CDS FW part 2 | CAAGACAAGGAGTTTACCTTGGAAGGACTATGAGATGGATGC |
| PHYA CDS RV part 2 | TCTTCGAGTGTGAAGACCATCTTGTTTGCTGCAGCGAGTT |
| NL3F10H FW | AACTCGCTGCAGCAAACAAGATGGTCTTCACACTCGAAGA |
| NL3F10H RV | CGATCGGGGAAATTCGAGCTTCAGTGATGGTGATGGTGAT |

**Note:** primers were designed with an overlap of 20bp corresponding to the ends of the destination vector (pGWB601) or adjacent DNA fragment.
